# Mechanochemical feedback and control of endocytosis and membrane tension

**DOI:** 10.1101/201509

**Authors:** Joseph Jose Thottacherry, Anita Joanna Kosmalska, Alberto Elosegui-Artola, Susav Pradhan, Sumit Sharma, Parvinder P. Singh, Marta C. Guadamillas, Natasha Chaudhary, Ram Vishwakarma, Xavier Trepat, Miguel A. del Pozo, Robert G. Parton, Pramod Pullarkat, Pere Roca-Cusachs, Satyajit Mayor

## Abstract

Plasma membrane tension is an important factor that regulates many key cellular processes. Membrane trafficking is tightly coupled to membrane tension and can modulate the latter by addition or removal of the membrane. However, the cellular pathway(s) involved in these processes are poorly understood. Here we find that, among a number of endocytic processes operating simultaneously at the cell surface, a dynamin and clathrin-independent pathway, the CLIC/GEEC (CG) pathway, is rapidly and specifically upregulated upon reduction of tension. On the other hand, inhibition of the CG pathway results in lower membrane tension, while up regulation significantly enhances membrane tension. We find that vinculin, a well-studied mechanotransducer, mediates the tension-dependent regulation of the CG pathway. Vinculin negatively regulates a key CG pathway regulator, GBF1, at the plasma membrane in a tension dependent manner. Thus, the CG pathway operates in a negative feedback loop with membrane tension which leads to a homeostatic regulation of membrane tension.

## Introduction

Living cells sense and use force for multiple functions like development^1^, differentiation^2^, gene expression^3^, migration^4^ and cancer progression^5^. Cells respond to changes in tension, passively by creating membrane invaginations/ blebs^6–8^ and actively, by modulating cytoskeletal-membrane connections, mechanosensitive channels and membrane trafficking^4,9,10^. Membrane trafficking through endo-exocytic processes can respond and modulate the membrane tension^10^. While exocytosis acts to reduce plasma membrane tension as a consequence of increasing net membrane area, endocytosis could function to reduce membrane area and enhance membrane tension.

Membrane tension has long been shown to affect the endocytic process. Decreasing tension upon stimulated secretion or by addition of amphiphilic compounds increases endocytosis ^11 12^. On the other hand, an increase in tension upon hypotonic shock ^11^ or as evinced during mitosis^12^, results in a decrease in endocytosis. Increase in membrane tension on spreading is also compensated via an increase in exocytosis from endocytic recycling compartment providing extra membrane^13^. Together these observations suggest that endocytosis responds to changes in membrane tension or changes in membrane area. However, the specific endocytic mechanisms involved in these responses have not been elucidated. The well-studied dynamin-dependent, clathrin-mediated endocytic (CME) pathway is relatively unaffected by changes in tension^14^ while caveolae help to passively buffer membrane tension^7^.

We had recently shown that upon relaxing the externally induced strain on cells, tubule like membrane invaginations termed as ‘reservoirs’ are created^6^. This initial response is a purely passive mechanical response of the plasma membrane and is also observed in synthetic lipid membranes. Similar structures are also formed on recovery from hypo-osmotic shock termed as ‘vacuole like dilations’ (VLDs). VLDs form passively, similar to reservoirs, albeit different in shape. Subsequent to the formation of either reservoirs or VLDs, cells engage in an active ATP-dependent response that occurs efficiently only at physiological temperatures to restore their morphology and membrane tension^6^. This indicates the deployment of specific active cellular processes following the passive response (see the cartoon in Fig 1a).

**Figure 1:**
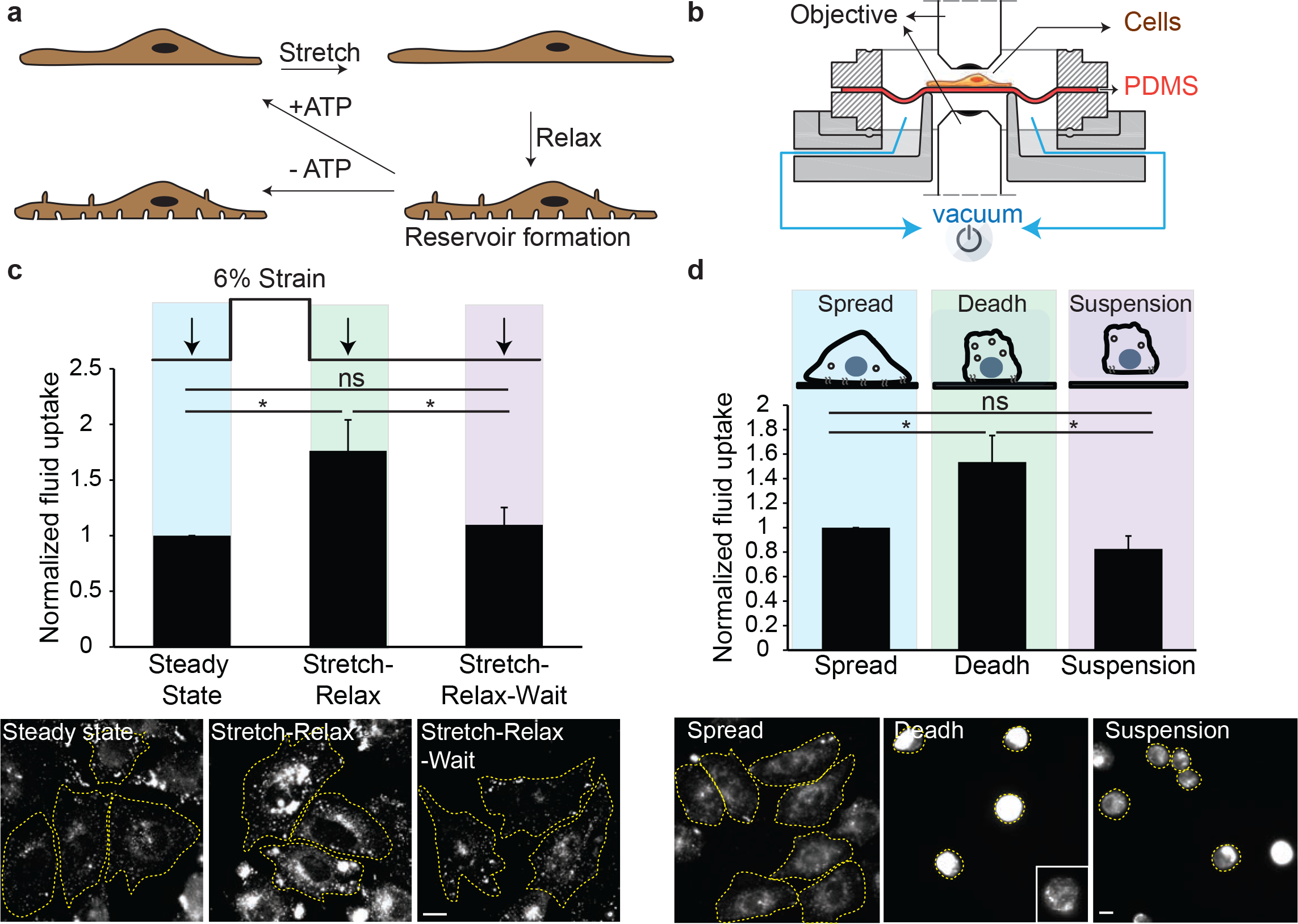
A fast transient endocytic response to decrease in membrane tension: **(a)** Cartoon showing membrane remodeling responses after mechanical strain. Cells after the stretch and relax protocol forms invaginations termed ‘reservoirs’^6^. These reservoirs are resorbed in few minutes by an active process and requires ATP. **(b)** The illustration shows the longitudinal section of a vacuum based equi-bi-axial stretching device. Cells plated on a PDMS sheet are stretched by the application of controlled vacuum below the circular PDMS sheet, which stretches it in a calibrated manner. Releasing the vacuum relaxes the strain on PDMS thus relaxing the cell. Cells plated on PDMS can be imaged in an upright or inverted microscope as required. **(c)** Endocytic response on strain relaxation. CHO cells were pulsed for 90 sec with TMR-Dex at steady state (steady state), immediately on relaxing the stretch (stretch-relax), or after a waiting time of 90 seconds after relaxing the stretch (stretch-relax-wait). After the pulse, cells were quickly washed with ice cold buffer and fixed, followed by imaging on a wide field microscope. Images show representative cells used to generate the histograms which provide a quantitative measure of the extent of endocytosis of TMR-Dex for the indicated treatments **(d)** Endocytic response on deadhering. CHO cells are pulsed with TMR-Dex for 3 minutes in adhered cells (Spread), during de-adhering (Deadh), or immediately after cells are detached and in suspension (Suspension) were washed with ice cold buffer, added back to concanavalin coated glass bottom dishes in ice cold buffer, fixed and imaged on a wide field microscope. Images and Histogram show the extent of fluid-phase uptake under the indicated conditions. In each experiment, the data represent the mean intensity per cell (± S.D) from two different experiments with duplicates containing at least 100 cells per experiment. *: *P* < 0.001, ns: not significant. Scale bar, 10 μm.

Here we have explored the nature of such active responses. We have tested the functioning of multiple endocytic pathways on modulation of membrane tension by different approaches. In parallel, we have determined the effects of modulating endocytic processes on membrane tension by utilizing optical or magnetic tweezers to measure membrane tension. Subsequent to the passive membrane response we had observed earlier^6^, we find that a clathrin, caveolin and dynamin-independent endocytic mechanism, the CLIC/GEEC (CG) pathway, rapidly responds to changes in membrane tension, acting to restore it to a specific set point. Perturbing the CG pathway directly modulates membrane tension forming a negative feedback loop with membrane tension to maintain homeostasis. A previously identified mechanical transducer, vinculin, is involved in the homeostatic control of tension; in its absence the CG pathway fails to respond to changes in membrane tension, thereby altering the set point.

## RESULTS

### A rapid endocytic response to changes in membrane tension

Active cellular processes are involved in resorbing the ‘reservoirs’ or ‘VLDs’ formed following a strain relaxation^6^. To determine whether endocytosis could be one such active process, we monitored the extent of endocytosis by providing a timed pulse of a fluid-phase marker, fluorescent-dextran (F-Dex), during and immediately after the stretch-relax procedure (using a custom built stretch device^6^ shown in Fig. 1b). Compared to cells at steady state, there was a dramatic increase in fluid-phase endocytosis immediately after relaxation of the strain (Fig. 1c) while uptake was markedly reduced during the application of the strain (Supplementary Fig. 1a). This increase in endocytosis was transient and disappeared as early as 90 seconds after strain relaxation (Fig. 1c). This also corresponds to the time scale of resorption of reservoirs by an active process observed earlier^6^. By rapidly upregulating endocytosis, cells thus respond to a net decrease in tension in a fast, transient fashion returning swiftly to a steady state.

Earlier studies indicated that exocytosis helped add membrane rapidly in response to increased membrane tension during cell spreading^15^. On deadhering, cells round off decreasing their surface area while on replating, cells spread by adding membrane. Thus it is likely that endocytic pathways could help retrieve membrane on deadhering due to decrease in net membrane tension^8,16^. We reasoned that if an endocytic process is responding to the release of strain during the deadhering process, it would be upregulated during the detachment process. To monitor the extent of endocytosis we followed the uptake of a timed pulse of F-Dex during and immediately after the detachment (3 minutes) and compared it to that measured in the spread state (Fig.1d schematic). Our results showed that the net fluid-phase uptake underwent a rapid increase while the cells were de-adhering (3 minutes), but subsided back to the steady state level once it was de-adhered and held in suspension (Fig. 1d). Recycling of the endocytic material is similar between steady state and deadhering (Supplementary Fig. 1b). This indicates that there is no block in the recycling rate during deadhering and increase in uptake on deadhering is due to a transient increase in endocytic potential.

To further consolidate our findings, we used an alternate method to alter membrane tension. We shifted cells from hypotonic to isotonic medium, which made passive invaginations similar to reservoirs called VLD’s^6^. This method also results in an enhancement of fluid-phase endocytosis (Supplementary Fig. 1c), consistent with the results obtained by the other two methods of altering membrane tension. Together these results suggest that reduction of membrane tension via a number of different methods triggered a fast and transient endocytic response on the time scale of seconds.

### Membrane tension and the response of multiple endocytic pathways

To ascertain which of the multiple endocytic pathways respond to changes in tension, we examined cargo previously shown to be endocytosed via these distinct pathways. A number of endocytic pathways function concurrently at the cell surface^17–20^. In addition to the well characterized CME pathway, there are pathways that are independent of clathrin but utilize dynamin for vesicle pinching^19,21^. Additionally, there are clathrin and dynamin independent pathways which function in a number of cell lines^22–24^, but not in all^25^. The CLIC/GEEC (clathrin independent carrier/ GPI-anchored protein enriched early endosomal compartment) pathway is a clathrin and dynamin-independent pathway, responsible for the internalization of a major fraction of the fluid-phase and several GPI-anchored proteins (GPI-AP)^22,24^, and other plasma membrane proteins such as CD44^26^. Therefore, we used these specific cargoes to test the response of different endocytic pathways while membrane tension was altered.

The endocytic uptake of the transferrin receptor (TfR), a marker of CME, did not increase in the cells which exhibited a transient rise in the fluid-phase after a hypotonic shock (Fig 2a) or detachment (Fig. 2b) as visualized using two color fluorescence microscopy. However, uptake of the folate receptor, a GPI-AP that is internalized via the CG pathway^27,28^, exhibited a considerable increase (Fig. 2c). This indicated that clathrin-independent endocytosis rather than CME might be involved in the fast response to a decrease in membrane tension.

**Figure 2:**
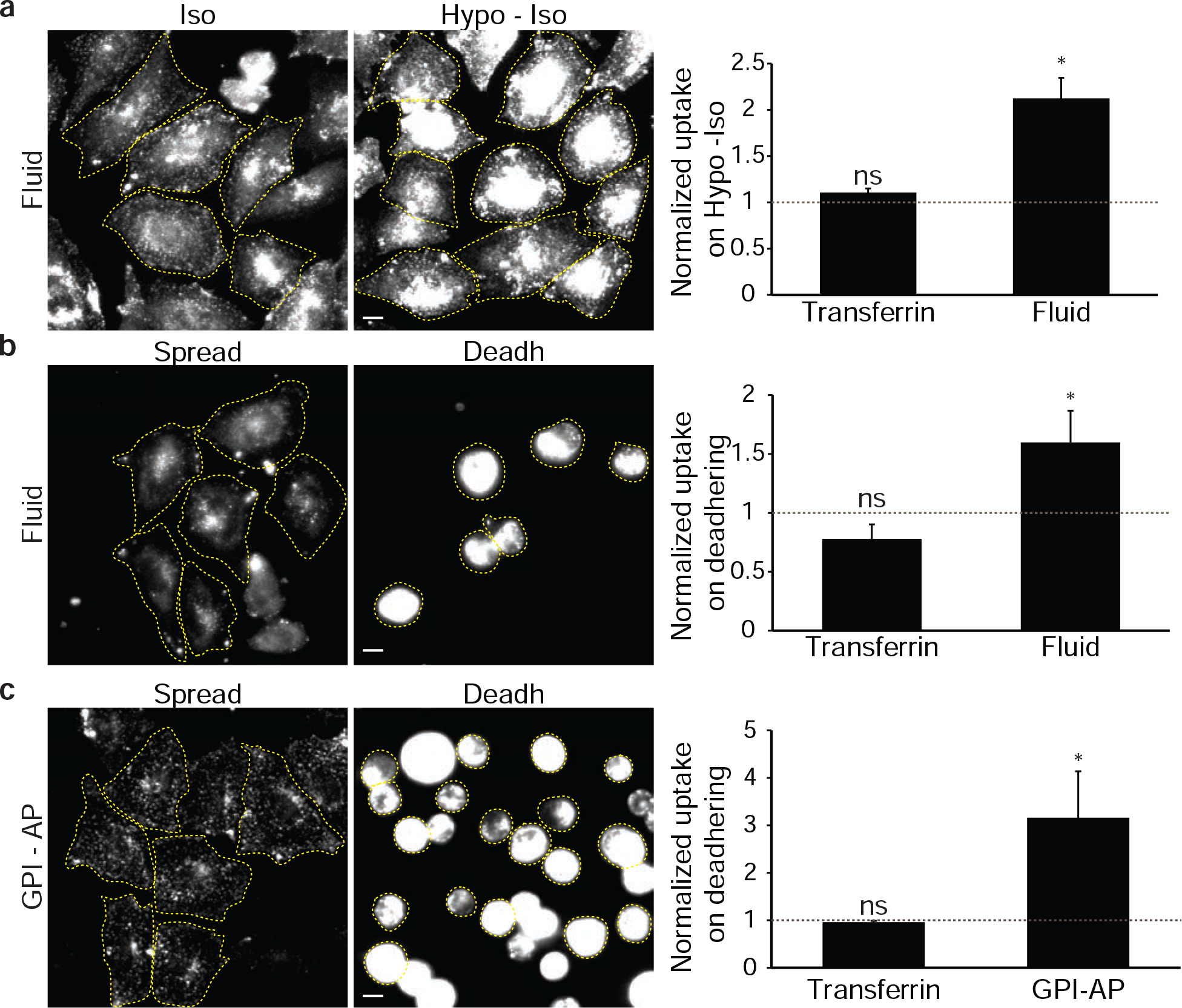
Endocytic pathways differ in their response to decrease in membrane tension: **(a)** Fluid-phase and transferrin (Tf) uptake in CHO cells on recovery from osmotic shock. Fluid-phase and Transferrin uptake were monitored in CHO cells under Isotonic conditions (Iso) or immediately after shifting from the hypotonic to isotonic state (Hypo-Iso) by incubating cells with A647-Tf (Transferrin) or TMR-Dex (Fluid) as indicated in Methods. Wide-field images (left) show the extent of endocytosed fluid-phase in Isotonic or Hypotonic-Isotonic (Hypo-Iso) conditions. Histogram (right) show the extent of TMR-Dex and A647-Tf endocytosis in the Hypo-Iso condition normalized to those measured in the isotonic condition (grey dashed line).**(b)** Fluid-phase and Tf uptake in CHO cells on deadhering. CHO cells were pulsed with TMR-Dex (Fluid) and A647-Tf (Transferrin) for 3 minutes when the cells are adherent (Spread) or during detachment (Deadh), and taken for imaging as described in Methods. Wide-field images (left) show the extent of endocytosed fluid-phase in Spread and during de-adhering condition (Deadh). Histogram (right) show the extent of TMR-Dex and A647-Tf endocytosis in the de-adhered condition normalized to that measured in the Spread condition (grey dashed line). **(c)** GPI-AP and Tf uptake on deadhering in CHO cells. CHO cells pulsed with fluorescent folate to label cell surface GPI-anchored folate receptors (GPI-AP) and with A647-Tf (Transferrin) for 3 minutes at 37 °C in normally adherent cells (Spread) or during detachment (Deadh), and taken for imaging as described in Methods. Wide-field images (left) show the extent of endocytosed GPI-anchored folate receptor in Spread and during the deadhering condition. Histogram (right) shows the extent of A647-Tf and Folate receptor endocytosis in the de-adhered condition normalized to those measured in the Spread condition (grey dashed line). In each experiment, the data represent the mean intensity per cell (± S.D) from two different experiments with duplicates containing at least 100 cells per experiment. *: *P* < 0.001, ns: not significant. Scale bar, 10 μm.

There are endocytic pathways which utilize dynamin independent of clathrin function^19,22^. Therefore we tested whether the increase in fluid-phase uptake requires dynamin function. We used a conditional triple knock out cell line that removes Dynamin 1, 2 and 3 from the genome^29^, thereby abolishing all the dynamin-mediated endocytic pathways (Supplementary Fig. 2a). The dynamin triple knockout mouse embryonic fibroblasts (MEFs) shows higher steady state fluid-phase endocytosis^29^. However, cells lacking all forms of dynamin also transiently increased their fluid-phase endocytosis upon both stretch-relax cycles to the same extent as wild type (WT)-MEFs (Fig. 3a) and hypotonic/isotonic media changes (Supplementary Fig. 2b). Thus, neither CME nor dynamin-dependent endocytic pathways appear to respond to an acute reduction in membrane tension.

**Figure 3:**
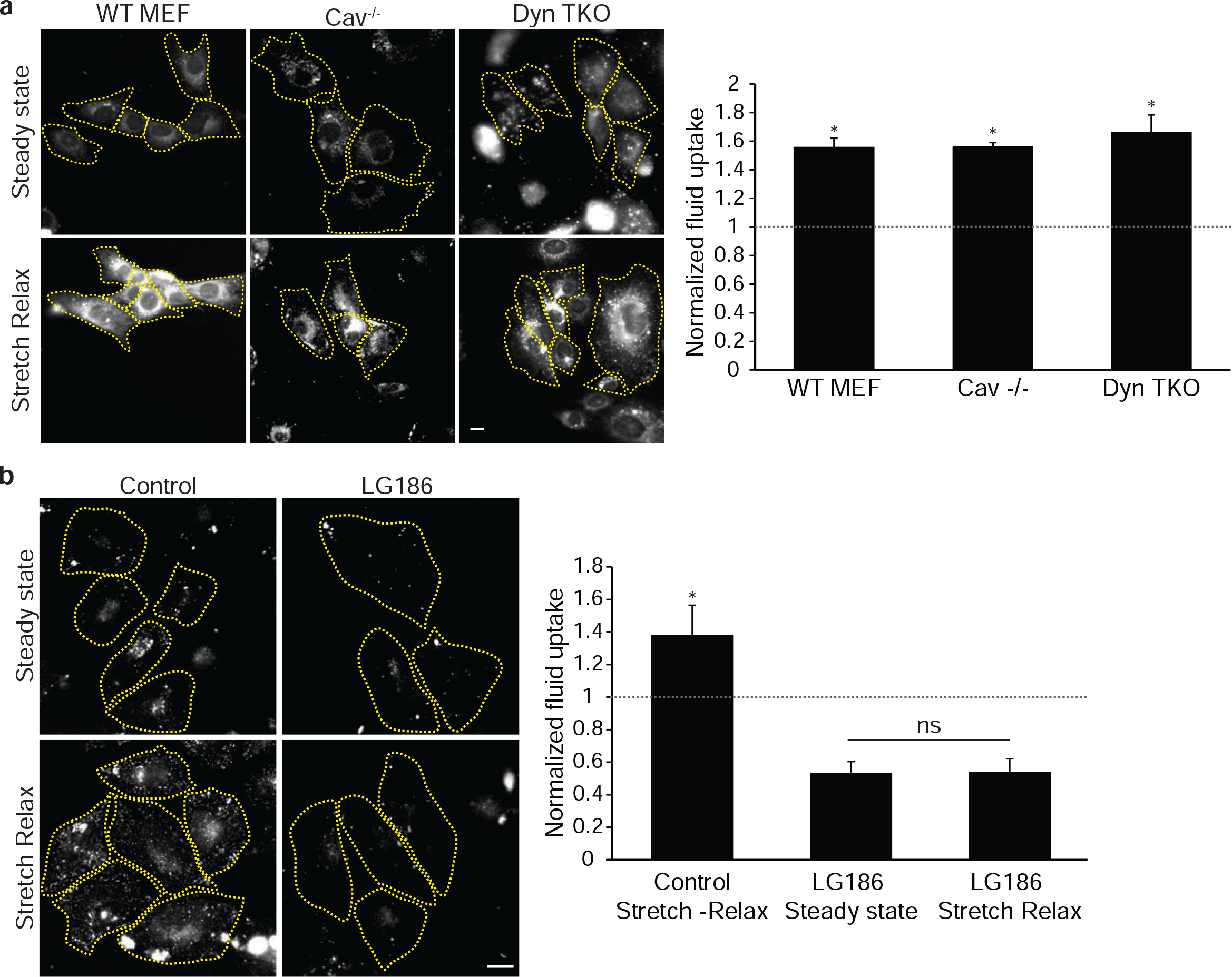
CLIC/GEEC (CG) pathway is the primary pathway for fast endocytic response: **(a)** Endocytic response in WT, dynamin TKO, and caveolin null MEFs on stretch-relax. Wild type MEF (WT MEF), Caveolin^-/-^ (Cav^-/-^), or conditional Dynamin triple knock out cells (Dyn TKO) were pulsed for 90 sec with TMR-Dex at steady state, immediately after relaxing the stretch (stretch-relax), and quickly washed with ice cold buffer, fixed and imaged on a wide field microscope. Images (left) show representative cells used to generate the histograms (right) which provide a quantitative measure of the extent of endocytosis of TMR-Dex for the indicated treatments. The uptake on stretch-relax in each cell line is plotted normalized to the steady state uptake in the respective cell lines (grey dashed line). **(b)** Inhibition of CG pathway and endocytosis on stretch-relax. CHO cells were treated with DMSO (Control) or with LG186 (10 μg/ml) to inhibit GBF1 for 30 minutes prior to pulsing with TMR-Dex for 90 sec, either at steady state (steady state), or immediately after relaxing the stretch (stretch-relax), and quickly washed with ice cold buffer, fixed and imaged on a wide field microscope. Images show representative cells used to generate the histograms (right) which provide a quantitative measure of the extent of endocytosis of TMR-Dex for the indicated treatments, normalized to the control steady state condition (grey dashed line). In each experiment, the data represent the mean intensity per cell (± S.D) from two different experiments with duplicates containing at least 100 cells per experiment. *: *P* < 0.001, ns: not significant. Scale bar, 10 μm.

A caveolin-dependent endocytic process is important to retrieve specialized membrane on deadhering^30^, and a caveolae-mediated passive mechanism is reported to buffer the increase in membrane tension and prevent cell lysis triggered by the flattening of caveolae^7^. To test if caveolin-dependent endocytic mechanisms could be important for this rapid endocytic up-regulation, caveolin null MEFs were subjected to the stretch-relax protocol. These cells exhibited a transient increase in fluid-phase uptake similar to their WT controls (Fig. 3a). In addition, caveolin-null cells also exhibit a fast transient upregulation of fluid-phase endocytosis during de-adhering as well (Supplementary Fig. 2c).

We next examined the morphology of the endocytic carriers formed by reduction of membrane tension induced by deadhering using electron microscopy (EM). For this, we utilized Cholera Toxin bound HRP (CTxBHRP), which marks the internalized plasma membrane. We used a procedure in which the surface remnant peroxidase reaction product is quenched with ascorbic acid, revealing only the internalized CTxBHRP labeled membrane^26^. After 5 minutes post-deadhering the major endocytic structures labeled had the typical morphology of CG carriers (or CLICs) comprising structures with tubular and ring-shaped morphology (arrows, Supplementary Fig. 2d). Morphologically-identical structures were also observed in WT MEFs at steady state^31^ and in Cav1^-/-^ MEFs (arrows, Supplementary Fig. 2d) consistent with the observation of fast fluid-phase uptake in Cav1^-/-^ cells via CG (Fig. 3a and Supplementary Fig. 2c). At this time point, surface-connected caveolae (containing no peroxidase-reaction product) persist in the Cav-expressing WT cells (arrowheads, Supplementary Fig. 2d), consistent with the possibility that the caveolar pathway does not play a significant role in transiently modulating endocytosis at these early times of deadhering.

Together, these experiments indicated that the clathrin, dynamin or caveolin dependent endocytic mechanisms do not exhibit a rapid respond to a reduction in membrane tension. This is in contrast to fluid-phase or GPI-anchored protein uptake which is endocytosed via the CG pathway. CG-mediated endocytosis is a high capacity pathway capable of internalizing the equivalent of the entire plasma membrane area in 12 minutes^26^, and of recycling a large fraction of endocytosed material^32^. This pathway is also implicated in the delivery of membrane on cell spreading in response to increases in membrane tension, thus helping to maintain membrane homeostasis^9,13^.

### CLIC/GEEC (CG) pathway responds to membrane tension

Since the CG cargo responded to changes in tension, we explored this finding in further detail. CG pathway is regulated by small GTPase’s, ARF1, its GEF GBF1, and CDC42 at the plasma membrane^28,33,34^. Hence, we utilized small molecule inhibitors of CDC42 and GBF1 to acutely inhibit CG pathway^35,36^. The CDC42 inhibitor, ML141 decreases fluid-phase endocytosis in cells at steady state but not CME (Supplementary Fig3a), and prevents the increase in fluid-phase uptake upon deadhering (Supplementary Fig. 3b). In separate experiments, we utilized LG186, an inhibitor of GBF1, which also decreases fluid-phase endocytosis in cells at steady state but does not affect CME (Supplementary Fig 3c). Inhibiting GBF1 prevents the increase in fluid-phase endocytosis observed upon stretch-relax (Fig. 3b) or deadhering (Supplementary Fig. 3d). Similar to the decrease in fluid-phase on increasing tension during stretch (Supplementary Fig. 1a), CD44, a CG pathway specific cargo, shows reduced endocytosis during hypotonic shock (Supplementary Fig. 3e).

To further confirm that this response is due to CG endocytosis, we assessed the effect of the stretch-relax protocol on cells that lack CG endocytosis. HeLa cells have been shown to lack a robust CG endocytic pathway^25,27^. While the molecular basis for this defect is not understood, we find that fluid-phase endocytosis in HeLa cells is not susceptible to GBF1 inhibition by LG186 (Supplementary Fig. 4a). In addition, these cells do not show an obvious recruitment of GBF1 to the plasma membrane in the form of punctae as observed in cells exhibiting constitutive CG endocytosis such as CHO cells as reported earlier^34^ (Supplementary Fig 4b/4c). Correspondingly, these cells did not exhibit a rapid increase in fluid-phase endocytosis on a hypotonic to isotonic shift (Supplementary Fig. 4d).

These experiments, combined with the up-regulated endocytosis of a CG specific cargo (GPI-AP) (Fig. 2c), suggest that the CG endocytic pathway is specifically involved in the rapid, transient response to changes in membrane tension.

### Passive and active membrane response to changes in membrane tension

As mentioned above, upon a rapid reduction in membrane tension, cells form passive structures such as reservoirs and VLDs similar to the response of an artificial membrane. Reservoirs are formed upon strain relaxation in the membrane after stretching cells, whereas VLDs are formed by water expelled by the cell after a hypo-to-isotonic-shock recovery^6^. Both reservoirs and VLDs are reabsorbed and disappear within a couple of minutes, coincidental with an increase in endocytosis. This led us to test if inhibiting the CG pathway could have a measurable impact on the rate of disappearance of such passive structures. We find that the CG pathway exhibits exquisite temperature sensitivity and is barely functional at room temperature (RT), and is not efficient even at 30°C in comparison to CME in CHO cells (Fig. 4a). Correspondingly, the reservoir resorption in CHO cells was impaired after lowering temperature (Fig. 4b). In addition, inhibition of the CG pathway in CHO cells by inhibiting GBF1 reduced the rate of reservoir reabsorption at 37 °C (Fig. 4a). In contrast, HeLa cells lacking a characteristic CG pathway did not show any difference in the rate of disappearance of reservoirs upon inhibiting GBF1 and was much less affected by lowering of temperature than CHO cells (Supplementary Fig. 4e). Thus, perturbing CG endocytosis affects the kinetics of resorption of reservoirs.

**Figure 4:**
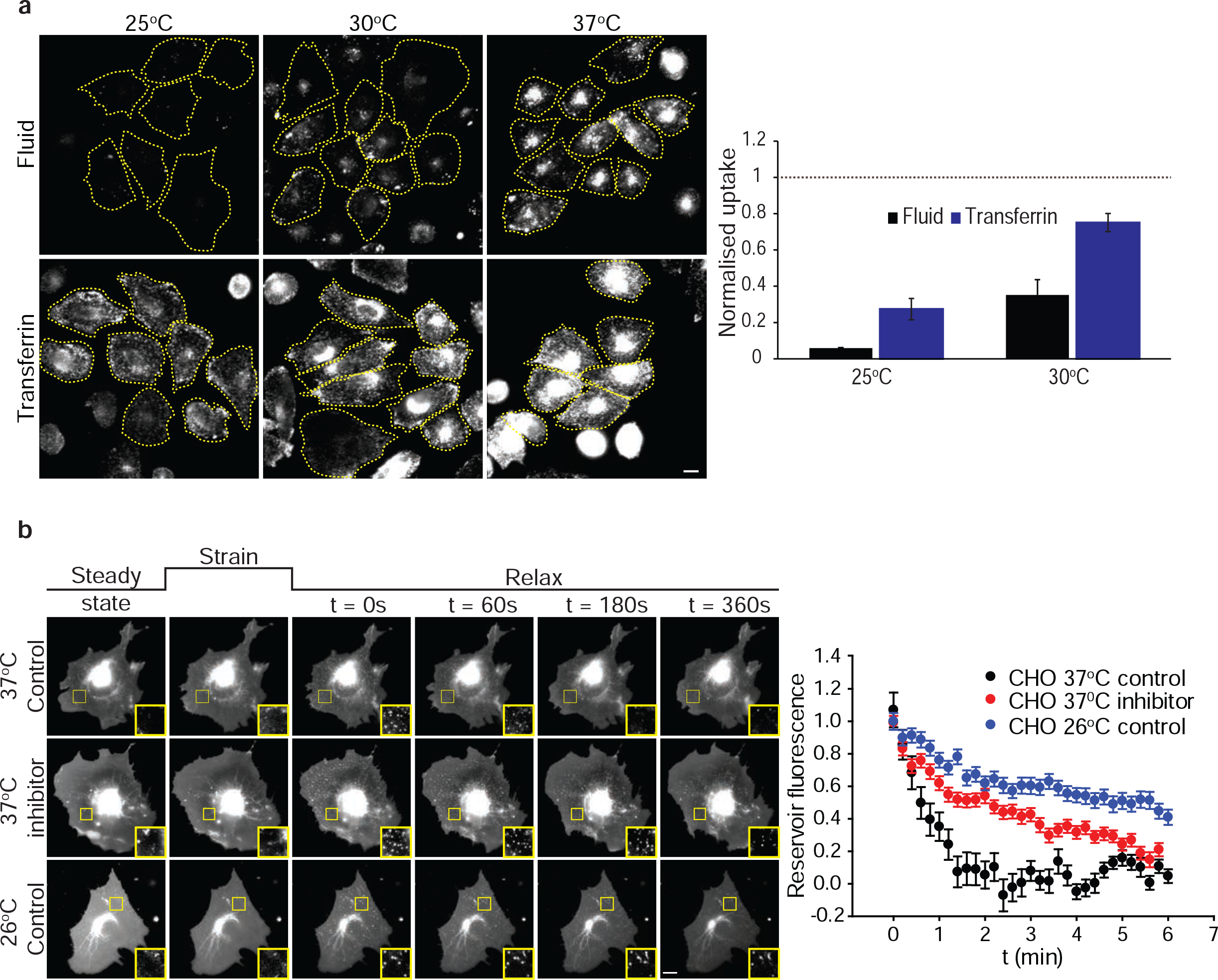
Temperature dependence of CG pathway and reservoir resorption: **(a)** Endocytosis of fluid-phase and Tf with temperature. CHO cells pre-equilibrated at the indicated temperatures were pulsed with TMR-Dex (Fluid) and Tf-A647 (Transferrin) for 5 minutes at the respective temperatures, and the extent of endocytosis of the two probes were quantified and normalized to those obtained at 37 °C (grey dashed line). Representative images (left) of cells used to generate the histogram (right) (mean± S.D) were obtained from two different experiments with duplicates each containing at least 100 cells per experiment. Scale bar, 10 μm. **(b)** Reservoir resorption on inhibiting CG pathway and decreasing temperature. The reservoir fluorescence intensity after stretch relax of CHO cells transfected with a fluorescent membrane marker (pEYFP-mem) was quantified as a function of time at 37°C in the absence (37°C control) or presence of LG186 (37°C inhibitor), or at room temperature (26°C control). Each point represents mean ± S.E.M from more than 100 reservoirs from at least 10 cells. Scale bar, 10 μm.

We next examined if passively generated structures could help initiate endocytosis at the sites of their formation. Since the disappearance of each reservoir is gradual and not as a single step process^6^ (Fig. 4b), this indicates that reservoirs are not likely to be pinched off directly as endosomes. Further, we do not observe endosomes form at the site of the reservoirs (Supplementary Fig. 5a). To test this, we took advantage of our earlier observation that cells plated on polyacrylamide gels do not form VLDs upon hypotonic to isotonic shifts^6^. Whereas the lack of generation of VLDs was confirmed in our cells grown on polyacrylamide (Supplementary Fig. 5b), the cells still showed an increase in endocytosis similar to when plated on glass, upon exposure to hypo-to-isotonic-shock procedure (Supplementary Fig. 5c). Together, these data suggest that CG endocytosis occurs subsequent to the passive responses of the membrane but the passive invagination formation is not necessary to form CG endosomes. However, the transient increase in CG endocytosis following the passive response helps swiftly resorb the excess membrane helping to restore the membrane morphology.

### Role of the CG pathway in setting membrane tension

Since the CG endocytic pathway responded to changes in membrane tension we hypothesized that it might be involved in the setting of steady state membrane tension as well. To explore this hypothesis, we directly measured tether forces by pulling membrane tethers using optical tweezers^37^. The force experienced by membrane tethers provides a way to measure the effective membrane tension^38^ (Fig 5a). We found that acutely inhibiting the CG pathway by inhibiting GBF1 drastically reduced the tether forces in a resting cell (Fig. 5b). To further assess this, we applied 0.5 nN force pulses using a magnetic tweezer device to ConcanavalinA (ConA)-coated magnetic beads attached to the cell membrane (Supplementary Fig 5d). Consistent with optical tweezers results, resistance to force (stiffness) was reduced in GBF1 inhibited cells (Supplementary Fig 5e). That this effect was due to a reduction in membrane tension and not any effects on the cytoskeleton, was corroborated by the lack of a change in the measured stiffness of fibronectin-coated beads attached to cells via integrin-fibronectin adhesions with and without GBF1 inhibition (Supplementary Fig 5f).

**Figure 5:**
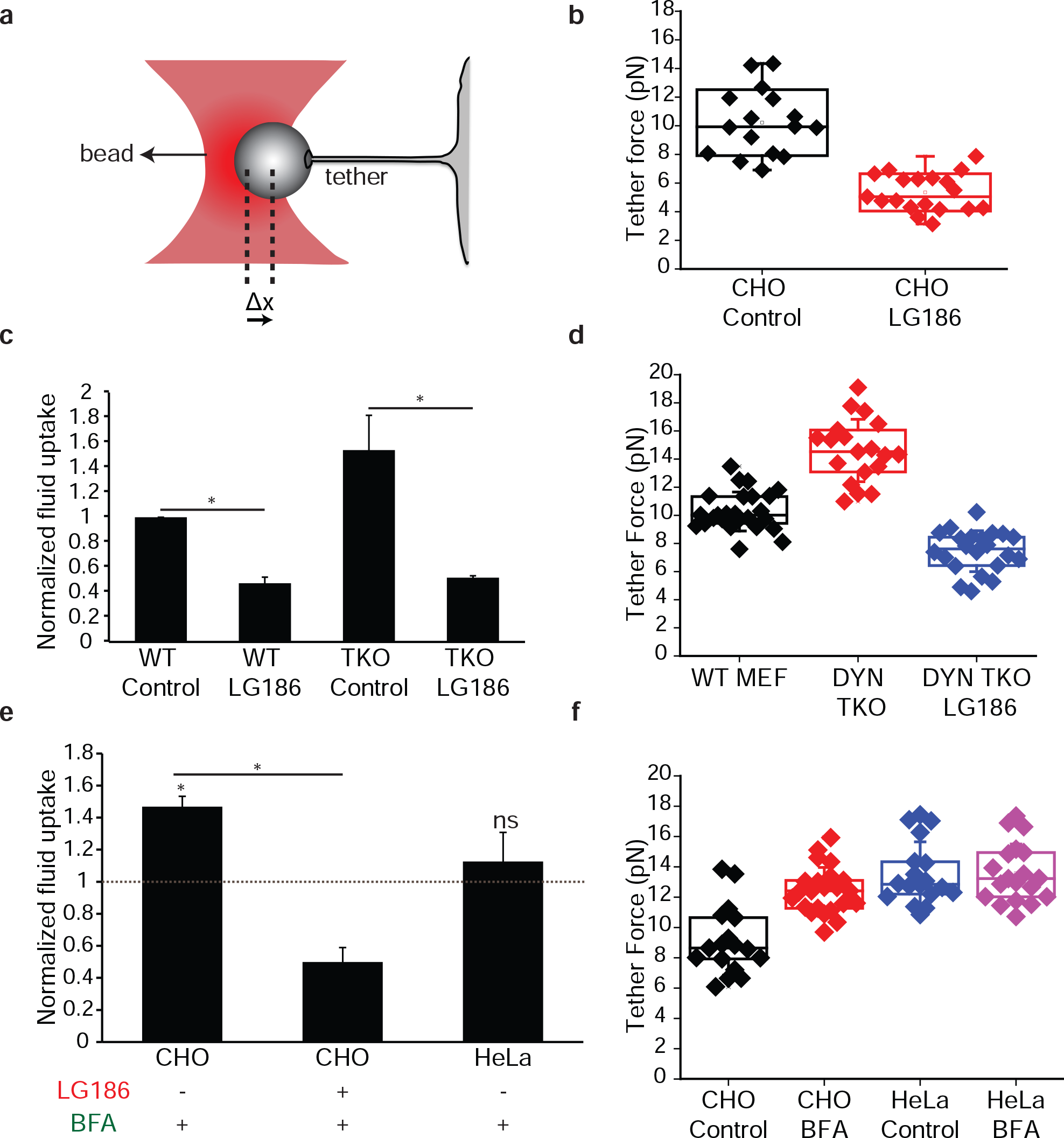
CG pathway regulates membrane tension: **(a)** Cartoon shows a membrane tether attached to a polystyrene bead trapped in an optical trap, used to measure tether forces. The polystyrene bead is held using a laser based optical trap to pull membrane tethers from cells. Displacement of the bead from the center of the trap (Δx) gives an estimate of the tether force (F) of the cell using the Hook’s law (F= −k*Δx; where k is the spring constant or trap stiffness; see methods). Membrane tension is obtained from the steady state force as σ = F^2^/(8π^2^B), where B is the bending modulus of the membrane and F is the tether force^58^. **(b)** Tether forces on downregulating CG pathway by GBF1 inhibition. Membrane tethers were pulled from CHO cells pre-treated with DMSO (CHO Control) or LG186 (CHO LG186) for 30 minutes, and maintained at 37 °C during the measurement. Tether forces were calculated as indicated above. The box plot shows data points with each point corresponding to a tether per cell with data combined (n = 16 (CHO control) and 19 (CHO LG186)) from two different experiments. **(c, d)** Endocytosis (c) and tether forces (d) in dynamin TKO cells. Wild type (WT) MEF, or conditional Dynamin TKO cells were either pre-treated with DMSO or LG186 and were pulsed for 5 minutes with TMR-Dex and taken for imaging (c) or directly taken to measure tether forces (d). The histograms shows fluid-phase uptake normalized to that observed in untreated WT MEF cells (c) and box plot shows tether forces (d) measured in the indicated conditions (n = 25 (WT MEF), 19 (DYN TKO) and 22 DYN TKO LG186)). **(e)** Endocytosis on modulating CG pathway. CHO cells were treated without BFA (Control), or with BFA (20μg/ml) alone or with LG186 for 30 minutes, and then directly incubated with TMR-Dex for 5 minutes, and imaged on a wide field microscope. The histogram shows fluid-phase uptake per cell normalized to control treated CHO cells (Dashed grey line). HeLa cells were correspondingly treated with BFA and the histogram shows the extent of fluid-phase uptake under the indicated conditions, normalized to that observed in untreated HeLa cells. **(f)** Box plot shows tether forces measured in CHO or HeLa cells treated without (Control) or with BFA for 45 minutes (n = 17 (CHO Control),23 (CHO BFA),18 (HeLa Control), 19 (HeLa BFA)). **(c, e)** In each endocytosis experiment, the data represent the mean intensity per cell (± S.D) from two different experiments with duplicates containing at least 100 cells per experiment. *: *P* < 0.001, ns: not significant.

We next examined tether forces in cells wherein the CG pathway is up-regulated. We reasoned that since the Dynamin TKO cells show a higher fluid-phase endocytosis (Fig. 5c, Supplementary Fig 6c), it is likely that this would increase effective membrane tension. Tether forces were indeed higher in the Dynamin TKO cells compared to control cells (Fig. 5d). Consistent with the role of the CG-pathway in setting membrane tension, inhibiting the CG pathway in Dynamin TKO cells by GBF1 inhibition (Fig. 5c, Supplementary Fig 5h) reduced the effective membrane tension below control levels (Fig. 5d).

To further confirm this observation, we measured tether forces on acutely increasing CG endocytosis by using BrefeldinA(BFA) as reported earlier^33^. BFA treatment disrupts ER to Golgi secretion but also serves to free up ARF1, making it available at the cell surface to increase CG endocytosis^33^. We further confirm that this increase is mediated through a GBF1-sensitive CG endocytosis (Fig 5e). We treated the cells with BFA and measured tether forces using optical tweezers when the increase in endocytosis was most prominent. Tether forces were higher on treating cells with BFA compared to the control case (Fig. 5f). BFA treatment inhibits secretion^39^ and this could also increase the effective membrane tension due to a reduction of membrane delivery from the secretory pathway independent of its effect on CG endocytosis. To test this, we treated HeLa cells with BFA. BFA treatment disrupted the Golgi in both CHO and HeLa cells (Supplementary Fig. 5g) consistent with its inhibition of the secretory pathway. However, neither fluid-phase uptake (Fig. 5e) nor the tether forces were affected in HeLa cells (Fig. 5f). This indicated that the increase in tension in CHO cells on BFA treatment is due to an increase in CG endocytosis and not due to a block in secretion in these timescales.

Hence, modulating the CG pathway by activating or inhibiting key regulators modifies effective membrane tension directly. Since CG pathway is negatively regulated by membrane tension, this indicates that CG pathway operates in a negative feedback loop with membrane tension. Since CG pathway is specifically modulated by tension, it is conceivable that the molecular machinery regulating CG pathway would be modulated by changes in tension.

### Mechanical manipulation of the CG endocytosis machinery

We tested if key regulatory molecules involved in different endocytic pathways could be directly modulated by changes in tension. GBF1 is involved in the CG pathway and re-localizes from the cytosol to distinct punctae at the plasma membrane upon activation as visualized using TIRF microscopy^33,34^. We imaged GBF1-GFP recruitment to the plasma membrane in live cells using TIRF microscopy, during a hypotonic shock and after recovering from it. GBF1 punctae were lost on hypotonic shock (Fig 6a and 6b) indicating a direct response by GBF1 on increasing tension. On the other hand, recovery from a hypotonic shock caused the rapid assembly of GBF1 punctae (Fig 6a and Fig 6b). In contrast, clathrin, which is localized from cytosol to membrane to help in CME is not affected by these similar changes in tension (Supplementary Fig 6a). These experiments indicated that molecular machinery involved in regulating the CG pathway was modulated by membrane tension unlike that of the CME pathway.

**Figure 6:**
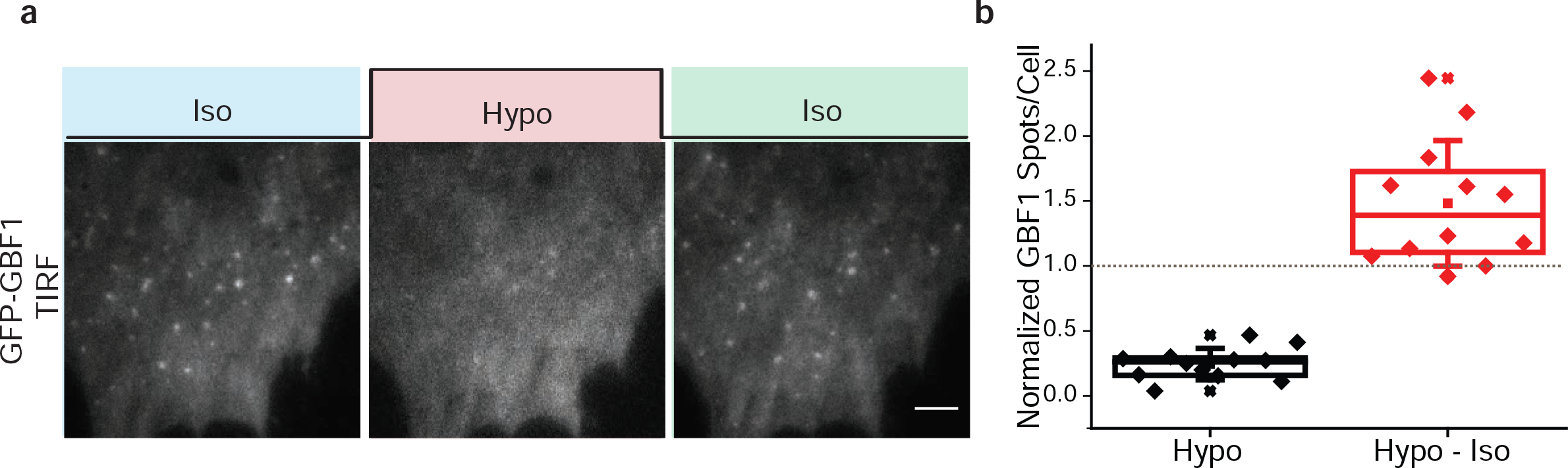
Mechanical modulation of CG molecular machinery. **(a)** WT- MEF cells transfected with GBF1-GFP were imaged live by TIRF microscopy. They exhibit GBF1-punctae at the plasma membrane, which is modulated by alterations in osmolarity obtained by changing the media from isotonic (Iso) to 40% hypotonic (Hypo) and back to isotonic (Iso). **(b)** Quantification of the number of punctae per cell during hypotonic shock and subsequent shift to isotonic medium. The GBF1 spots upon hypotonic shock and subsequent shift to isotonic medium is normalized to number of spots in the respective cell determined before hypotonic shift and plotted as a box plot. Each data point is a measurement from a single cell and box plot shows data of 12 cells from two independent experiments. Scale bar, 10 μm.

### Vinculin serves as a mechanotransducer for CG endocytosis

For cells to respond to changes in tension, sensing and transduction of this information must occur. Since focal adhesion related molecules help transduce and respond to force^5,40,41^ we hypothesized these molecules could transduce a physical stimuli (membrane tension) to regulate endocytic processes as well. Indeed, few of these proteins were ‘hits’ in a recent RNAi screen for genes that influence CG endocytosis^42^. Focal adhesion is an intricate macromolecular complex that has multiple functional modules^43^ and vinculin is a critical part of this mechanotransduction machinery^40,41,43^. Unlike the ‘hits’, Talin or p130CAS, there is only a single functional isoform of vinculin in non-muscle cells, and cell lines deficient for this protein are viable. Therefore we used a vinculin null fibroblast cell line to test the role of this mechanotransducer in the mechano-responsive behavior of CG pathway.

**Figure 7:**
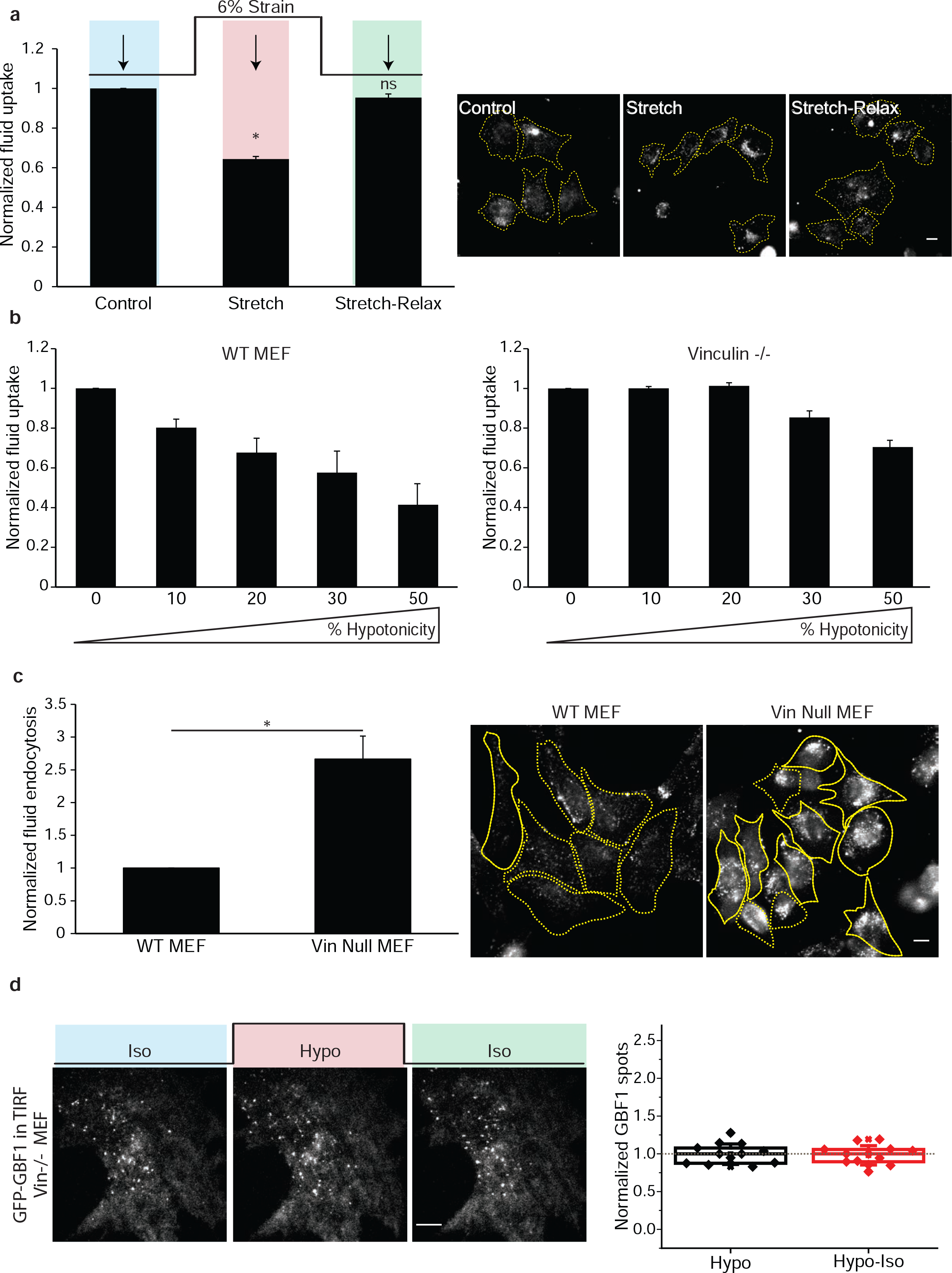
Vinculin dependent mechanoregulation of CG pathway. **(a)** Vinculin null cells were pulsed for 90 seconds with F-Dex either during a 6% stretch or on relaxing this strain. Fluid-phase uptake per cell during this strain change is normalized to uptake in steady state cells for the same time point, and plotted in the left panel. Representative images are shown in the right panel. **(b)** WT MEF (left panel) and vinculin null cells (right panel) are pulsed with F-Dex for 2 minutes in increasing hypotonic medium as indicated, washed, fixed and imaged. Average uptake per cell in the indicated hypotonic media was normalized to the average uptake of the isotonic condition and plotted as a box plot. Note while fluid-phase uptake is inhibited in WT MEF cells proportional to the increase in hypotonicity, vinculin null cells are refractory to much higher hypotonicity. **(c)** WT or vinculin null MEFs are pulsed with F-Dex for 3 minutes as described before at 37°C. Left panel shows quantification of uptake per cell normalized to the WT levels and right panel shows the representative images of the same. In each endocytosis experiment, the data represent the mean intensity per cell (± S.D) from 2 independent experiments with each with two technical duplicates containing at least 100 cell per experiment. *: *P* < 0.001, ns: not significant. Scale bar, 10 μm. **(d)** Vinculin null cells transfected with GBF1-GFP and imaged live using TIRF microscopy while changing media from isotonic (Iso) to 40% hypotonic (Hypo) and back to isotonic (Iso). GBF1 organization at plasma membrane during the osmotic shifts is shown in a representative cell (left panel). Quantification of the number of punctae per cell during hypotonic and isotonic shifts is normalized to number of spots (Grey dotted line) before hypotonic shift of the respective cells is plotted as a box plot (right panel). Each data point is measurement from a single cell and box plot shows data of 13 cells from two independent experiments. Scale bar, 10 μm.

To directly test if vinculin could be involved in the tension-sensitive regulation of the CG pathway, we stretched vinculin null cells. Unlike WT MEFs that shows ~82% drop in uptake on stretching (Supplementary Fig 1a), vinculin null cells show only ~36% drop at the same strain (Fig 7a). Increasing the extent of hypotonic shock showed a concomitant decrease in fluid-phase endocytosis of WT cells (Fig 7b). By contrast, vinculin null MEFs were much more refractory to the same extent of hypotonic shock (Fig 7b). Furthermore, upon strain-relaxation, fluid-phase endocytosis in vinculin null MEFs did not show an increase (Fig 7a), unlike that observed for CHO cells (Fig 1c) or wild type MEFs (Fig 3a). This was further tested in the deadhering assay where vinculin null cells again did not show an increased fluid-phase uptake, unlike the WT control cells (Supplementary Fig 6c). Thus vinculin null cells do not respond to changes in tension similar to the WT cells.

Fluid-phase uptake in vinculin null MEFs is much higher than wild type cells (Fig 7c). To test if the endocytic effects of vinculin null cells are specifically due to vinculin, we expressed full length vinculin in vinculin null cells. This caused a decrease in fluid-phase endocytosis (Supplementary Fig 6d). Further, inhibiting GBF1 in vinculin-null cells with LG186 decreased fluid-phase uptake to the same levels as cells expressing vinculin, confirming that a GBF1 sensitive CG pathway is functional here (Supplementary Fig 6d). This indicates that GBF1 operates downstream of vinculin and vinculin negatively regulates CG pathway.

Since vinculin-null cells have a higher basal endocytosis rate it is possible that they are unable to increase their endocytic capacity further in response to a decrease in membrane tension. However, vinculin null cells respond to BFA treatment to increase their endocytic rate in a manner that is also sensitive to GBF1 inhibition, similar to wild type cells (Supplementary Fig 7a).

We next tested if GBF1 shows a tension-dependent membrane localization of GBF1 in vinculin null cells. The level of punctae remained constant and failed to respond to hypotonic shock (Fig 7d) unlike that observed in WT MEF (Fig 6a). The density of GBF1 punctae at the plasma membrane was also slightly higher in vinculin null cells compared to WT cells (Supplementary Fig 6b). This is consistent with higher fluid-phase endocytosis in vinculin null cells compared to control MEF cell line (Fig 7c).

Further, we tested how the steady state membrane tension in vinculin null cells is compared to WT cells. Tether forces measured using optical tweezers showed a higher value for vinculin null cells compared to wild type cells (Fig 8a). The high tether force in cells lacking vinculin was drastically reduced on inhibiting the CG pathway (Fig 8a), consistent with the role of the CG pathway in regulating the effective membrane tension. These experiments show that vinculin acts as a negative regulator of CG pathway and is necessary for the transduction of physical stimuli for the biochemical control of the CG pathway.

**Figure 8:**
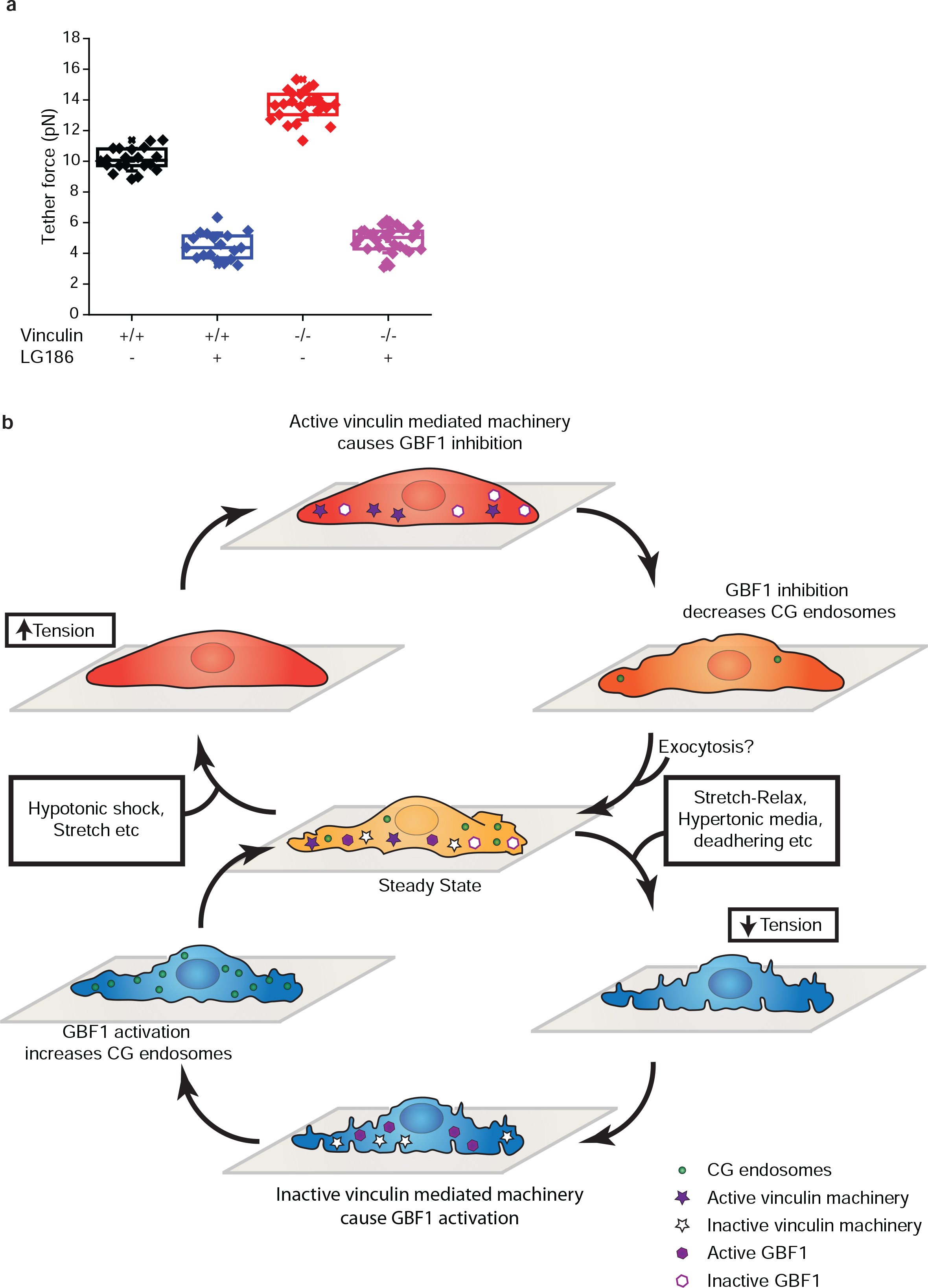
Membrane tension and Vinculin. **(a)** WT (Vin +/+) or vinculin null cells (Vin −/−) were treated with LG186 to inhibit GBF1 mediated CG pathway and membrane tension measured using optical tweezer as described before and compared to the control treated cells. (n = 20 (Vin +/+), 25 (Vin +/+ with LG186), 25 (Vin −/−) and 29 (Vin −/− with LG186)). Vinculin null cells shows a higher basal membrane tension compared to WT MEF; inhibiting the CG pathway drastically reduced membrane tension in both cell lines. **(b)** CG pathway and membrane tension operates in a vinculin dependent negative feedback loop to maintain homeostasis. Reduction of tension from its steady state leads to a passive response by the formation of reservoirs or VLDs. The decrease in the effective tension inactivates a vinculin-dependent machinery, resulting in an increase in active GBF1, which increases the CG pathway and rapid internalization of the excess membrane. This is a fast transient response that appears to restore the steady state. On the other hand, increasing the membrane tension from steady state activates vinculin dependent machinery, inhibiting the CG pathway, via the reduction of GBF1 recruitment. The increase in effective tension could also activate exocytic machinery which adds membrane resulting in restoration of the steady state. Thus, a vinculin dependent mechanochemical-regulation of the CG pathway through a negative feedback loop helps in maintaining plasma membrane tension homeostasis.

## DISCUSSION

Membrane tension has been long proposed to be tightly coupled to vesicular trafficking through endo-exocytic pathways. However, the specific trafficking mechanisms have remained elusive. Here, we show that membrane tension and CG endocytosis operate in a negative feedback loop that helps restore any change from a set point (model: Supplementary Fig. 8b). When membrane tension decreases, it transiently triggers the CG pathway, bringing about a fast endocytic response to reset the cell’s resting membrane tension. On the other hand, increasing membrane tension has the opposite effect; the CG endocytosis is inhibited in a proportional manner. The membrane flux through the CG pathway also has an effect on the effective membrane tension. Acutely lowering the CG pathway decreases membrane tension while upregulating the pathway increases membrane tension. Thus, changes in membrane tension lead to an inverse effect on CG endocytosis, while changes in CG endocytosis lead to a direct effect on plasma membrane tension. This type of a response known as a negative feedback loop is used in many different biological contexts to maintain homeostasis^44^.

The CG endocytic pathway specifically responds to acute changes in membrane tension despite multiple pathways operating simultaneously at the plasma membrane. Caveolae passively buffer increases in tension^7^, while the clathrin-mediated pathway concentrates specific ligands and mediates robust endocytosis despite the increase in tension^14^. De-adhered cells exhibit an increased caveolin-mediated internalization that persists over hours, and is crucial for the removal of specific membrane constituents and anchorage-dependent growth and anoikis^30^. On the other hand, unlike the caveolar pathway, the CG pathway showed a higher transient upregulation of endocytosis only during deadhering which does not persist in suspension.

The fast response of the CG pathway on strain relaxation is lost within 90 seconds. The loss of this transient response could be even faster as at present 90 seconds is the dead time in our experiments. Fast clathrin-independent mechanisms have been reported in different contexts. Ultrafast endocytosis occurs at synapses following a synaptic vesicle fusion to retrieve the excess membrane. This fast clathrin-independent but dynamin-dependent process is temperature sensitive as well^45,46^. Endophilin dependent FEME pathway is another clathrin-independent but dynamin-dependent pathway. HeLa cells seem to predominantly have AP2, GRAF1 and dynamin-dependent machinery^25^. Consistent with this, inhibiting AP2 or GRAF1 in HeLa cells inhibits fluid-phase endocytosis. These cells respond in a slower rate to the changes in osmotic shock and shows blebbing on inhibiting this pathway due to lack of endocytosis^47^. Here, we find that the GBF1/ARF1/CDC42 dependent CG pathway shows a fast transient response to changes in tension and is more sensitive to lowering of physiological temperature compared to the CME pathway. In the absence of such a fast pathway, other slower endocytic mechanisms operate to internalize the excess membrane, lack of which might lead to blebbing. The CG pathway helps to swiftly respond and reset any changes from the steady state, thereby also helping to set the resting membrane tension of a cell. This indicates that different endocytic pathways have distinct functions and the CG pathway may be responsible for membrane tension homeostasis.

Similar to the endocytic response, exocytic processes in a cell could modulate and respond to changes in tension. Exocytic processes in a cell help in addition of membrane to the cell surface and reduction of membrane tension^9^. Unlike endocytosis, exocytosis seems to be positively regulated by membrane tension. High membrane tension increases the exocytic rate and could regulate the mechanism of vesicle fusion^48,49^. Increase in membrane tension during cell spreading activates exocytosis to increase spread area through a GPI-anchored protein rich endocytic recycling compartment^13,15^. This increase in area is independent of secretory pathway or other exocytic mechanisms. CG pathway takes in a major fraction of GPI-anchored proteins^27^ and recycles a huge fraction of its endocytic volume^32^. On increasing tension, we find that CG endocytosis is downregulated preventing further increase in tension but it could be recycling through CG pathway that helps add membrane to restore the steady state tension (Cartoon: Fig 8b). Thus, regulation of membrane cycling through the endo-exocytic leg of CG pathway could be important in membrane homeostasis and further research would help bring about a complete view of this mechanism.

We find that the active response by CG pathway follows the passive response via membrane invaginations (i.e. reservoirs and VLD) and helps in efficient resorption of these passive local membrane structures. There could be other active cellular mechanisms driving the flattening of these invaginations as well. However, these membrane invaginations are not necessary for the creation of the CG endosomes. Thus, following a reduction in tension these are two parallel responses by the cell, one passive and the other active eventually leading to excess membrane internalization through CG endosomes.

Similar to the passive response, physical parameters could directly regulate the active endocytic machinery by influencing the extent of membrane deformation needed to make an endocytic vesicle. A higher membrane tension makes it more difficult to deform the membrane, thus producing fewer endosomes, and vice versa, alleviating the need for a specific mechanotransduction machinery. However, our results from studying the vinculin-null cells suggest otherwise. Vinculin, a key focal adhesion protein, transduces many mechanical inputs at the site of the focal adhesion into information for the cell to process^41,43,50^. In this context, it appears that vinculin plays a central role in transducing the increase (or decrease) in membrane tension to the CG pathway to help inhibit (or activate) its endocytic mechanism (Cartoon: Fig 8b). This appears to be effected by its control of a key regulator of CG endocytosis, GBF1, the GEF for ARF1. In WT cells, GBF1 forms tension-sensitive punctae at the cell surface wherein increasing tension abolishes these punctae and decreasing tension increases it. In the vinculin-null cells, this tension-dependent regulation is lost and CG endocytosis appears to be uncoupled from tension regulation. Thus, vinculin is important for a tension-sensitive negative regulation of a key effector of CG pathway, translating mechanical information into a biochemical read out to influence the endocytic rate. This negative feedback loop between effective membrane tension and CG pathway thus is mediated through vinculin and maintains the cells at a lower effective membrane tension. Different functional modules operate in a focal adhesion for mechanotransduction^43^. However, the precise mechanism behind the ability for vinculin to regulate the availability of GBF1 at the cell surface is not yet understood and is a subject of further investigation.

Modulations in membrane tension are used in multiple cellular processes^4,51–54^ and CG pathway could have a role in these. In migrating fibroblasts, CG endosomes are localized to the leading edge and transient ablation of these endosomes inhibits efficient migration^26^. Increase in the membrane tension at the leading edge keeps the neutrophil cells polarized and helps in its migration^54^. In a separate study in neutrophils, GBF1 localizes to the leading edge by binding to products of phosphatidylinositol 3-kinase (PI3K), recruits ARF1, and this localization is needed for unified cell polarity^55^. We have found that PI3K products help recruit GBF1 to the plasma membrane and this is necessary for CG endocytosis^56^. Thus, one could speculate that a polarized CG pathway and its regulators operating in migrating cells could be modulating membrane tension by regulating membrane trafficking.

Endocytic pathways are proposed to be at the core of a eukaryotic cellular plan integrating multiple inputs over spatio-temporal scales^57^. They are also necessary for tissue patterning: the CG pathway is utilized for Wingless signaling for patterning of the *Drosophila* wing disc during larval development^56^. Here we find that a high capacity CG pathway that turns over a huge fraction of the plasma membrane^26,32^ and sensitive to membrane composition^32^ is modulated by temperature and mechanochemical inputs. A vinculin-mediated negative feedback loop between membrane tension and the CG pathway helps maintain the cell at a lower tension set point (Cartoon: Fig 8b). This could also help in increasing the potential for modulating membrane tension to regulate other cellular processes. Thus, the CG pathway responds and coordinates a variety of cellular inputs including membrane tension and is likely to function in multiple physiological contexts.

## ACKNOWLEDGEMENTS

We thank Pietro De Camilli (Yale University, USA) for conditional Dynamin triple knock out cell line, Daniel Rösel (Charles University, Prague) for vinculin null cell line, Darius V Köster for the Caveolin null cell line, David J Stephens (University of Bristol, UK) for an initial gift of LG186, Feroz M.H. Musthafa (CCAMP, Bangalore), G.V. Soni (RRI, Bangalore) for help with preparation of PDMS membrane. We would like to thank Manoj Mathew and central imaging and flow cytometry facility (CIFF, NCBS) for help with imaging, Dev Kumar (Mech. Workshop) for making components for stretch relax apparatus and imaging, Dr. Anusuya Banerjee for help with illustrations, K. Joseph Mathew for final cartoon (Fig 8b), and thank members of P.P, X.T, and P.R-C laboratories for hosting and helping J.J.T with day-to-day experiments. X.T. acknowledges support from the Spanish Ministry of Economy and Competitiveness (BFU2015-65074-P), the Generalitat de Catalunya (2014-SGR-927), and the European Research Council (ERC-2013-CoG-616480). This study was also supported by grants SAF2014-51876-R from Spanish Ministry of Economy and Competitiveness (MINECO) and co-funded by FEDER funds to M.A.dP, and 674/C/2013 from Fundació La Marató de TV3 to P.R-C and M.A.dP. R.G.P. was supported by the National Health and Medical Research Council (NHMRC) of Australia (program grant, APP1037320 and Senior Principal Research Fellowship, 569452), and the Australian Research Council Centre of Excellence (CE140100036). We acknowledge the Australian Microscopy & Microanalysis Research Facility at the Center for Microscopy and Microanalysis at The University of Queensland. J.J.T acknowledges pre-doctoral fellowship from Council for Scientific and Industrial Research (CSIR), Government of India. S.M would like to acknowledge J.C. Bose Fellowship from DST, Government of India and Wellcome-DBT Margdarshi fellowship.

## AUTHOR CONTRIBUTIONS

J.J.T and S.M conceived the study. J.J.T, A.J.K, P.P, P.R-C, M.A.d.P, R.G.P and S.M designed the experiments. J.J.T, A.J.K, A.E-A and N.C performed the experiments and analyzed them. M.C.G and R.G.P performed the EM experiments designed by R.G.P and M.A.d.P. S.P and P.P built the optical tweezer setup. X.T and P.R-C designed and built the stretch system. S.S, P.P.S and R.V synthesized LG186. J.J.T and S.M wrote the paper.

## SUPPLEMENTARY FIGURE LEGENDS

**Supplementary Figure 1:**
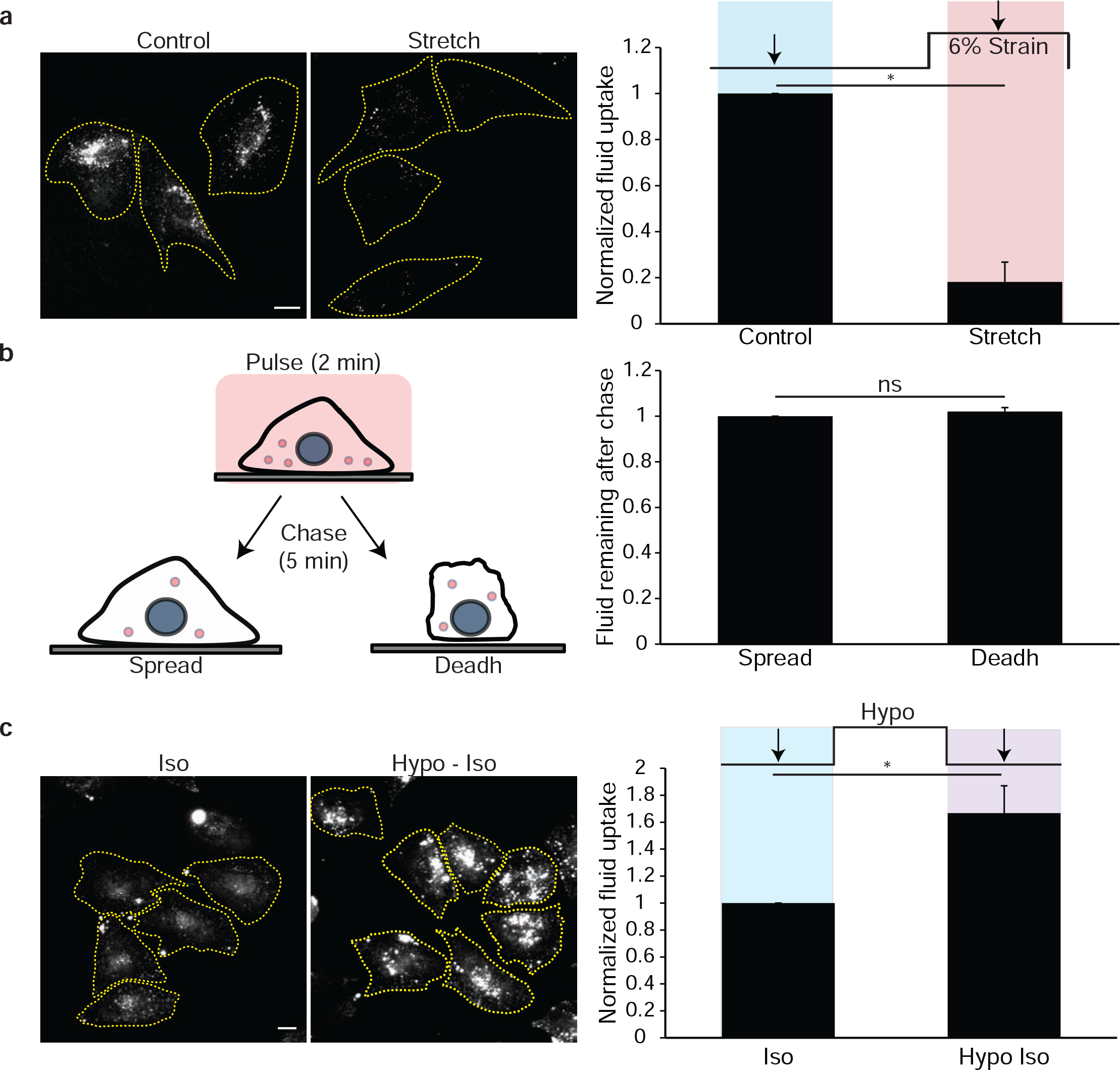
**(a)** Fluid-phase endocytosis is inhibited on stretching of cells. WT – MEF cells grown on the PDMS device as detailed in Methods, were subjected to 6% strain and pulsed for 90 seconds with TMR-Dextran (TMR-Dex) under the stretched condition (stretch), fixed and imaged. Fluorescence intensity is compared to the cells that were not subjected to the strain (control) cells pulsed for 90 seconds. **(b)** Recycling of the fluid-phase is not affected during deadhering. Cartoon describes the pulse chase protocol used to look at recycling during deadhering. CHO cells are pulsed with TMR-Dex for 2 minutes, washed and chased for 5 minutes either in their adhered steady state (Spread) or during de-adhering (Deadh) were washed, fixed, and imaged. Histogram (right) shows TMR-Dex remaining in the cells after chase in the deadhering or in spread condition normalized to the spread condition. **(c)** Shifting from hypotonic to isotonic media results in an increase in fluid-phase uptake. Cells were pulsed with TMR-Dex for 1 minute either in steady state (Iso) or after hypotonic shock (hypo-iso) for one minute. Images (left) of endocytosed TMR-Dex and histogram (right) show the extent of fluid-phase uptake. In each experiment, the data represent the mean intensity per cell (± S.D) from two different experiments with duplicates containing at least 100 cells per experiment. *: *P* < 0.001, ns: not significant. Scale bar, 10 μm.

**Supplementary Figure 2:**
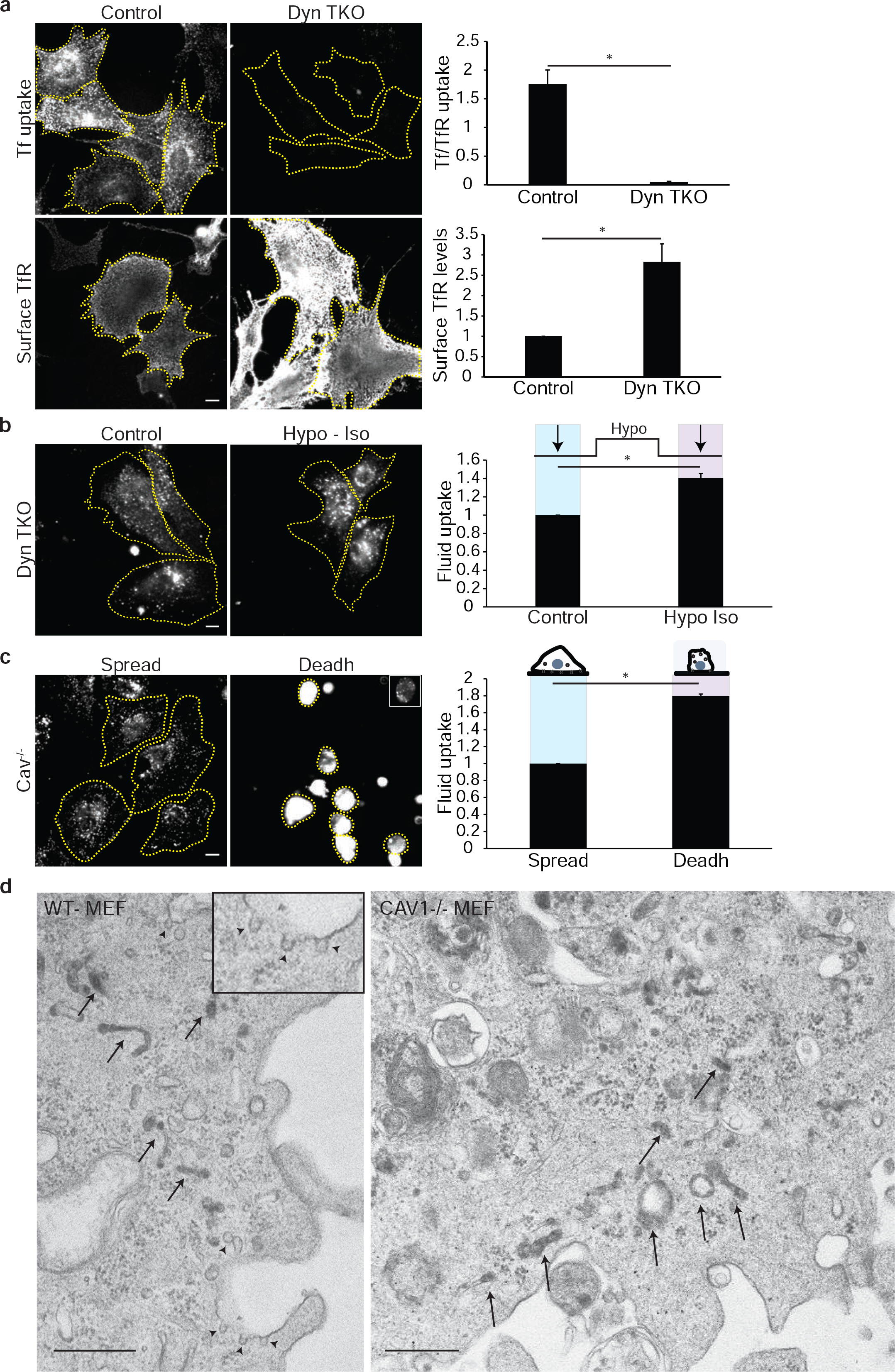
**(a)** Dynamin TKO cells as detailed in Methods (Dyn TKO), or (Control) was pulsed with Tf-A568 for 5minute, surface stripped, labelled with antibody against transferrin receptor (TfR), washed and fixed. Transferrin (Tf) uptake normalized to the TfR levels is plotted in the histogram (right top) along with the surface TfR normalized to the control (right bottom), determined per cell. Corresponding wide field images show extent of endocytosed TfR (top) and surface TfR (bottom). Note that removal of all Dynamin isoforms inhibits transferrin uptake comprehensively while increasing surface levels of TfR. **(b)** Dynamin TKO cells (as in panel a) were pulsed with TMR-Dex for 1 minute either in steady state (control) or after hypotonic shock (hypo-iso). Wide-field images (left) show extent of TMR-Dex uptake and the histogram (right) shows fluid-phase uptake per cell in hypo-iso condition normalized to the control. Note while uptake of TfR is completely inhibited (a), fluid-phase uptake (b) exhibits a typical increase on shifting from hypotonic to isotonic conditions in Dynamin TKO cells. **(c)** MEFs (Cav−/−) were pulsed with TMR-Dex for 3 minutes when they are normally adherent cells (Spread) or during detachment (Deadh). Wide field images and histogram show the extent of fluid-phase uptake under these conditions. **(d)** Electron micrographs of WT-MEF and Cav1−/− MEF on deadhering show similar CG endosomes. CTxB-HRP uptake for 5 minutes and processed for DAB reaction as described in methods is done in both wild type MEF (WT-MEF) and Cav1−/− MEF. Arrows show internalized CG carriers in both cell types and arrow head show surface connected caveolae in de-adhered WT cells (inset shows zoomed in image). In each experiment, the data represent mean intensity per cell (± S.D) from two different experiments each with technical duplicates containing at least 100 cells per experiment. *: *P* < 0.001. Scale bar, 10 μm **(a, b, c)**, 500nm **(d)**.

**Supplementary Figure 3:**
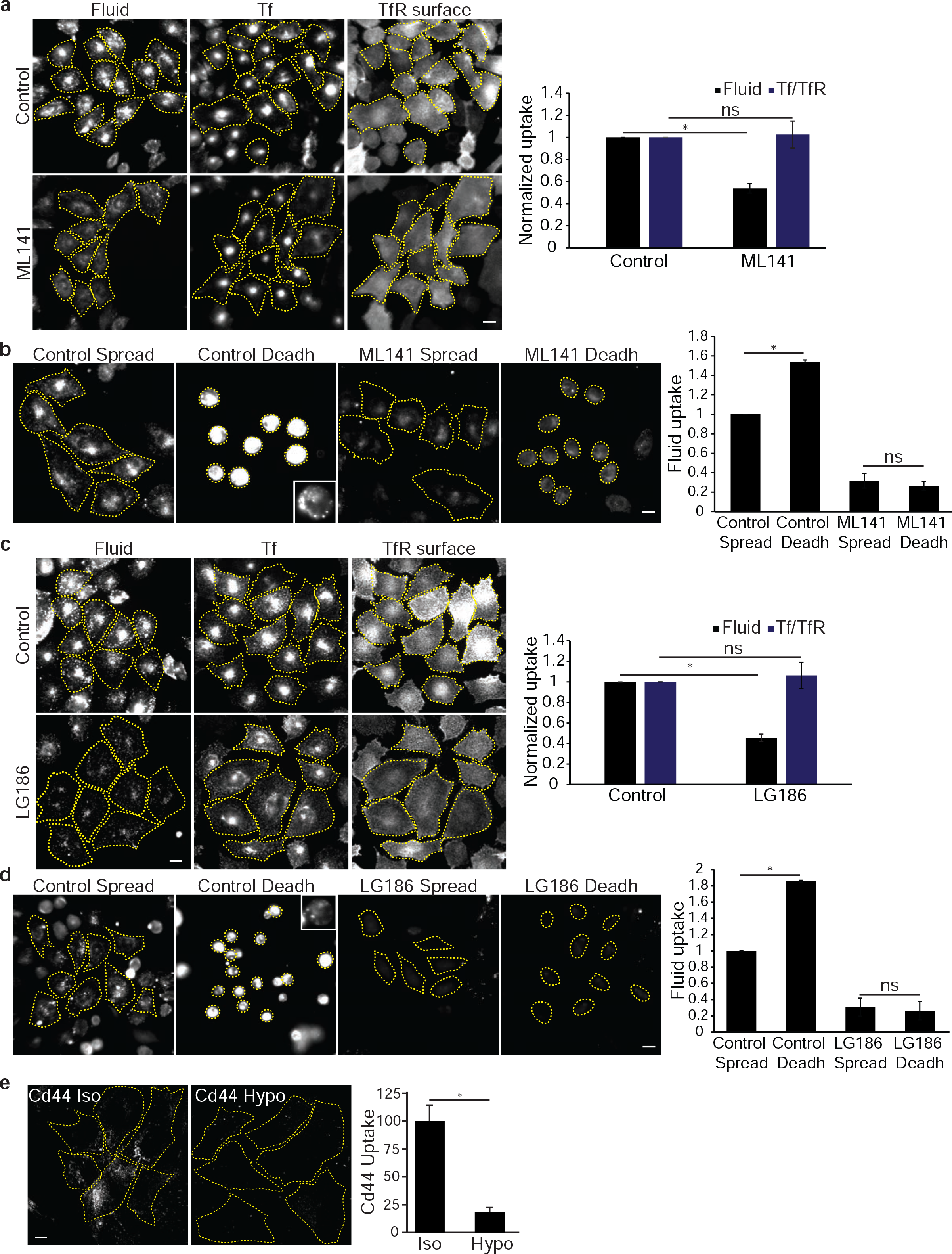
**(a)** ML141, a small molecule inhibitor of CDC42, inhibits fluid-phase uptake but not transferrin uptake at steady state. CHO cells treated without (Control) or with ML141 (10μM) for 45 minutes were pulsed for 5 minutes with TMR-Dex (Fluid) and A488-Tf (Tf), washed and surface-stripped of remnant-cell surface Tf. These cells were incubated with A647-labelled OKT9 antibody on ice to detect surface transferrin receptor (TfR Surface). Wide field images of cells (left) and the histogram of total fluid-phase and Tf uptake normalized to surface receptor level shows that the effect of ML141 was only on the fluid-phase uptake but not on TfR endocytosis. **(b)** Inhibiting CG pathway using ML141 prevents increase in fluid-phase uptake on deadhering. Adherent CHO cells treated without (Control) or with ML141 (10μM) for 45 minutes were pulsed for 5 minutes with TMR-Dex without detachment (Spread) or during the deadhering process (Deadh), and washed extensively, fixed and taken for imaging. Images (left) of endocytosed TMR-Dex and histogram (right) show the extent of fluid-phase uptake normalized to that observed in the control spread cells. Note that increase in endocytosis observed while deadhering is completely abolished upon inhibition of CG endocytosis by ML141. **(c)** LG186, a small molecule inhibitor of GBF1, inhibits fluid-phase uptake but not transferrin uptake. CHO cells treated without (Control) or with LG186 (10μM) for 30 minutes were pulsed for 5 minutes with TMR-Dex (Fluid) and A488-Tf (Tf), washed extensively and surface-stripped of remnant-cell surface Tf. These cells were incubated with A647-labelled OKT9 antibody to detect surface transferrin receptor (TfR Surface). Wide field images of cells (left) and histogram of total fluid-phase uptake and Tf uptake normalized to surface receptor level shows that the effect of LG186 was only on the fluid-phase uptake but not on TfR endocytosis. **(d)** Inhibiting CG pathway using LG186 prevents increase in fluid-phase uptake on deadhering. Adherent CHO cells treated without (Control) or with LG186 (10μM) for 30 minutes were pulsed for 5 minutes with TMR-Dex without detachment (Spread) or during the deadhering process (Deadh), and washed, fixed and taken for imaging. Images (left) of endocytosed TMR-Dex and histogram show the extent of fluid-phase normalised to that observed in the control cells. Note that increase in endocytosis observed while deadhering is completely abolished upon inhibition of CG endocytosis by inhibiting GBF1 using LG186. In each experiment, the data represent the mean intensity per cell (± S.D) from two different experiments with duplicates containing at least 100 cells per experiment. **(e)** CD44 uptake is inhibited at high tension. WT MEFs were pulsed CD44 mAb for 2 minutes either in isotonic medium (Iso) or in 75mOsm hypo-osmotic medium (Hypo) at 37°C, acid washed to strip surface mAb, washed and labeled with AF-555 secondary antibody. (Histogram (right) shows per cell uptake in hypo-osmotic situation normalized to the isotonic situation for the representative images shown on left. Data represents mean± S.E.M from three independent experiments).*: *P* < 0.001, ns: not significant. Scale bar, 10 μm.

**Supplementary Figure 4:**
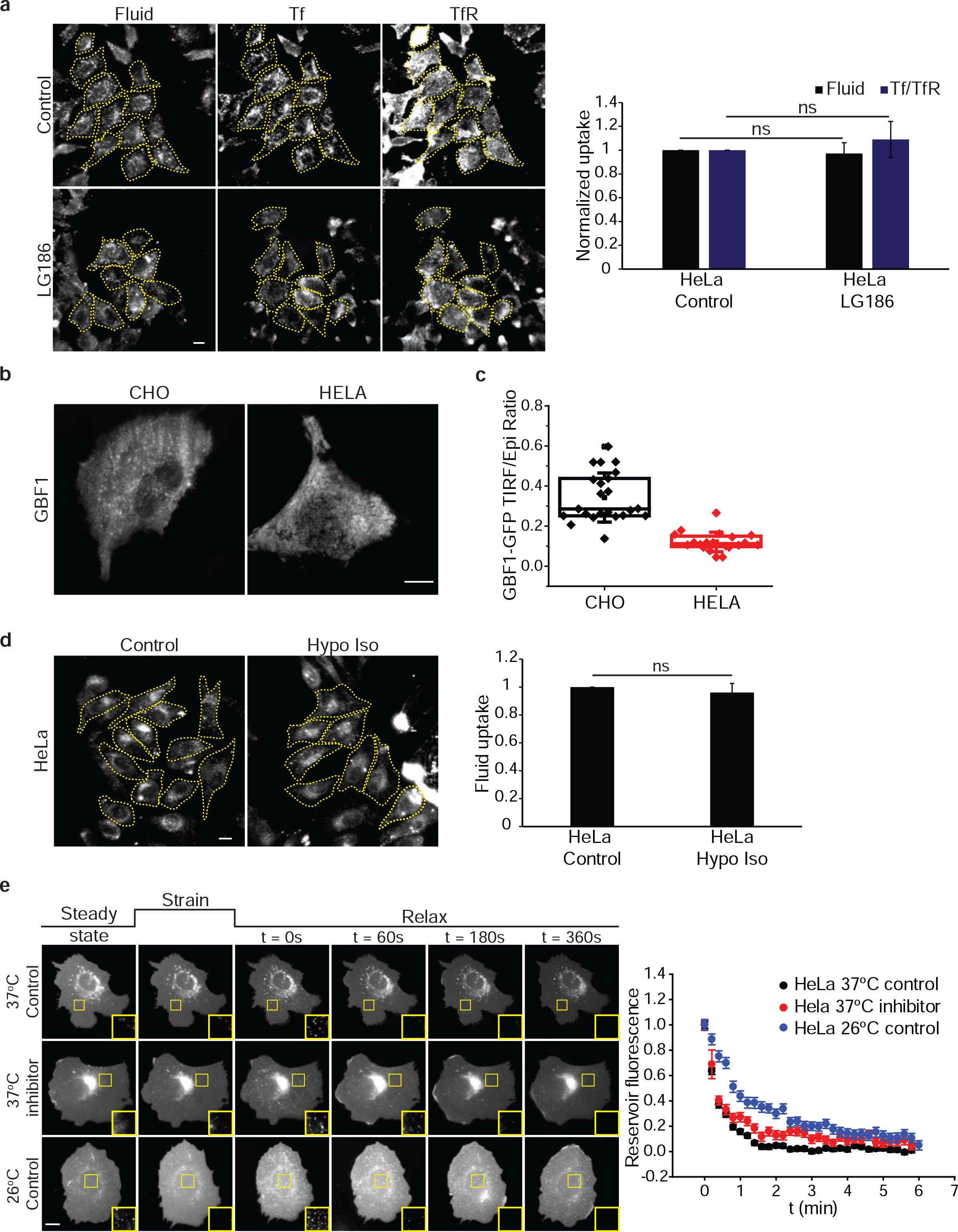
**(a)** Fluid-phase uptake in HeLa cells is insensitive to GBF1 inhibition by LG186 treatment. HeLa cells treated without (Control) or with LG186 (10μM) for 30 minutes were pulsed for 5 minutes with TMR-Dex (Fluid) and washed extensively, or pulsed with A488-Tf (Tf) and surface-stripped of remnant-cell surface Tf, followed by labeling with A648-labelled OKT9 antibody to detect surface transferrin receptor (TfR Surface). Wide field images of cells (left) and histogram of total fluid-phase uptake and Tf uptake normalized to surface receptor level shows that LG186 does not have an effect on the fluid-phase uptake and TfR endocytosis in HeLa cells. Data represent the mean intensity per cell (± S.D) from two different experiments with duplicates containing at least 100 cells per experiment. Scale bar, 10 μm. **(b**/**c)** GBF1 do not efficiently localize to the plasma membrane nor form punctae. GBF1-GFP was transfected in either CHO or HeLa cells and imaged in TIRF and in wide field (epi-fluorescence) format. HeLa cells do not show surface punctae formation unlike CHO cells (b). Quantification of the TIRF to wide field ratio of GBF1-GFP levels shows (c) that GBF1 translocation to surface is much lower in HeLa cells. Each point is value from a cell and quantified from two different experiments (n = 23 (CHO) and 20 (HeLa)). **(d)** Rapid shifting from hypotonic to isotonic state does not result in an increase in fluid-phase uptake in HeLa Cells. HeLa cells were pulsed with TMR-Dextran (fluid) for 1 minute either in steady state (Iso) or after hypotonic shock of one minute (Hypo-Iso). Images (left) of endocytosed TMR-Dex and histogram show the extent of fluid-phase uptake. Data represent the mean intensity per cell (± S.D) from two different experiments with duplicates containing at least 100 cells per experiment. **(e)** Reservoir adsorption in HeLa cells does not respond to GBF1 inhibition. The reservoir fluorescence intensity after stretch relax of HeLa cells transfected with a fluorescent membrane marker (pEYFP-mem) was quantified as a function of time at 37°C in the absence (37°C control) or presence of LG186 (37°C inhibitor), or at room temperature (26°C control). Each point represents mean ± s.e.m from more than 200 reservoirs from at least 10 cells. Note while treatment with LG186 had no effect on the rate of reservoir resorption, lowering of the temperature also had only a marginal reduction in the rate of resorption. Scale bar, 10 μm.

**Supplementary Figure 5:**
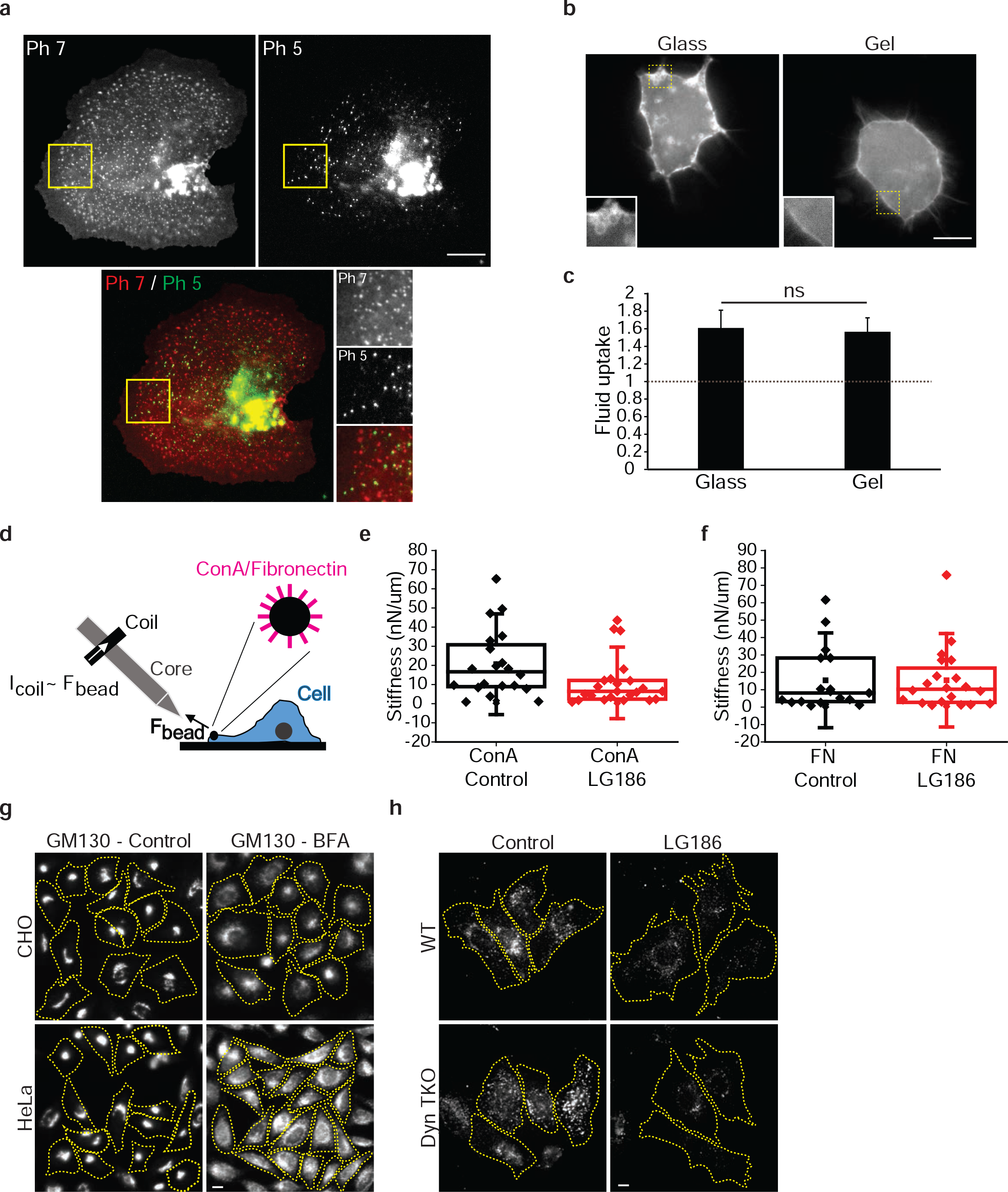
**(a)** Reservoirs formed on stretch-relax does not colocalize with endosomes. A pH sensitive SecGFP-GPI transfected CHO cells are stretched for one minute and relaxed in pH 7.4 buffer (pH 7, top left) at 37°C showing formation of reservoirs as before. The pH is made acidic (pH 5.5) after 30 seconds to quench surface fluorescence. Only internal vesicles at neutral pH remain fluorescent and thus helps to capture newly formed endosomes on stretch-relax (pH 5, top right). The reservoirs does not colocalize with the newly formed endosomes (bottom, insets). **(b)** The cells transfected with CAAX-GFP to mark the membrane are plated on glass (left) or polyacrylamide gels. Cells were incubated with hypo-osmotic medium for one minute followed by isotonic recovery. Images of VLDs are observed only in the cells grown on glass. Insets show a magnification of the edge of the cells where VLDs are expected to form. **(c)** Endocytic pulse of 90 seconds is done either in isotonic situation or after release of hypotonic shock of 1 minute in cells plated on glass or gel. Mean uptake per cell after release of hypotonic shock for either gel or glass is normalized to its uptake in isotonic conditions (dotted line). Data represent the mean intensity per cell (± S.D) from two different experiments with duplicates containing at least 100 cells per experiment. **(d)** Cartoon depicts a magnetic tweezer set up to measure membrane stiffness. An electromagnet with a sharpened tip was used to apply a 0.5 nN pulsatory force (1 Hz) to paramagnetic beads coated with either fibronectin fragment (FN) or ConcanavalinA (ConA), attached to cells as described in methods. Bead movement in response to applied force was then tracked, and bead stiffness was calculated during the first 10 seconds of the measurement in units of nN/μM. **(e, f)** Inhibiting of CG pathway with LG186 lowers membrane stiffness. ConA (e) or Fibronectin fragment (f) coated paramagnetic beads were attached to cells treated without (Control) or with LG186 for 30 minute. Beads were pulled using the magnetic tweezer as described, and membrane stiffness calculated. Note LG186 treatment lowers membrane stiffness only when ConA-coated beads are pulled, whereas FN-coated beads are unaffected. This indicates that the LG186 treatment does not disrupt the cortical cytoskeleton that interacts with the integrin-engaged FN-coated beads, but lowers general membrane tension as experienced by the ConA-coated beads. The box plot shows the stiffness values in nN/μm in each of the tethers pulled for the different conditions. **(g)** BrefeldinA (BFA) treatment causes disruption in GM130 distribution in both HeLa cells and CHO cells. Cells were treated with 20 μg/ml of BFA for 45 minute, fixed, permeabilized and labelled with GM130 antibody to mark cis-Golgi and imaged on a wide field microscope. BFA treatment disrupts the peri-nuclear localization in both CHO and HeLa cells, indicating that BFA is functional in both cell types. **(h)** The increase in fluid-phase uptake in Dynamin TKO is inhibited by LG186 treatment. WT or Dynamin TKO cells (prepared as described in methods) were treated with LG186 or with the vehicle (DMSO; Control), and incubated with TMR-Dex for 5 minutes, washed and fixed prior to imaging on a Wide Field microscope. The images show that while Dyn TKO cells exhibit a higher fluid-phase uptake that the WT cells, this uptake is sensitive to the inhibition of GBF1, confirming that it is the CG endocytosis that has a higher activity in Dynamin TKO cells (quantified in Fig. 5c). Scale bar, 10 μm.

**Supplementary Figure 6:**
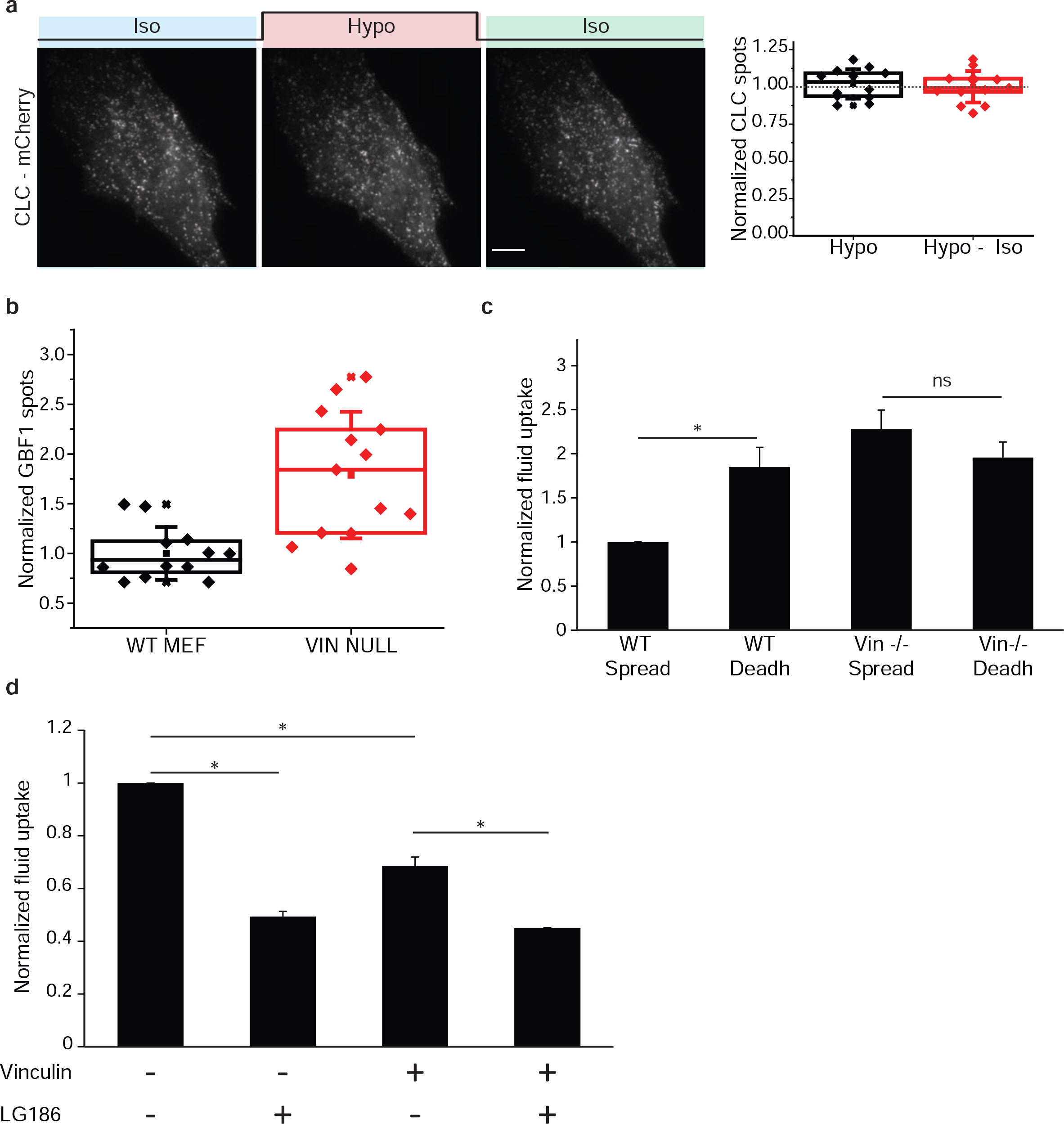
**(a)** WT cells transfected with clathrin light chain (CLC-mCherry) was imaged live using TIRF microscopy during osmotic changes as indicated. Images (Left panel) and graph (right) show the quantification of number of CLC punctae per cell normalized to the steady state number of punctae from 13 cells. Note that the number of clathrin pits at the cell surface do not respond to changes in tension. Scale bar, 10 μm. **(b)** GBF1 punctae in vinculin null cells are higher. GBF1-GFP is transfected in either WT-MEF or vinculin null MEF and imaged in TIRF. The number of punctae per cell were counted and normalized to the surface area. The plot shows the average number of spots in each cell type normalized to those obtained in WT-MEF. Each point in the box plot represents measurements collected from a single cell (WT MEF, n= 12; (VIN NULL, n= 13), accumulated. **(c)** Cells were pulsed with F-Dex either during deadhering or in spread state for 3 minutes and the pulse is stopped using ice cold buffer as described before. De-adhered cells are added to vial containing ice cold buffer and allowed to attach on a fresh coverslip bottom dish to be fixed and imaged. Endosomal intensity per cell is quantified and normalized to the intensity of the WT spread. Note that Vinculin null cells show higher endocytosis compared to WT cells but does not show increase in uptake on deadhering unlike WT cells. **(d)** Vinculin null cells rescued by transfection with WT-Vinculin full length (VIN+) and vinculin null cells were treated with LG186 to check sensitivity to GBF1 inhibition of fluid-phase endocytosis. Cells treated with LG186 for 30 minutes were pulsed for 5 minutes with F-Dex, washed, fixed and imaged. Endocytic uptake per cell is quantified and after normalizing to the untreated Vin null case (Control) is plotted as shown. Note adding back vinculin reduces the fluid-phase uptake. However in both in the Vinculin null as well as in the cells in which Vinculin is added back, endocytic uptake of the fluid-phase remains sensitive to LG186, indicating a role for vinculin in negatively regulating the CG pathway. Data represent the mean intensity per cell (± S.D) from two different experiments with technical duplicates with at least 100 cells per experiment. *: *P* < 0.001, ns: not significant.

**Supplementary Figure 7:**
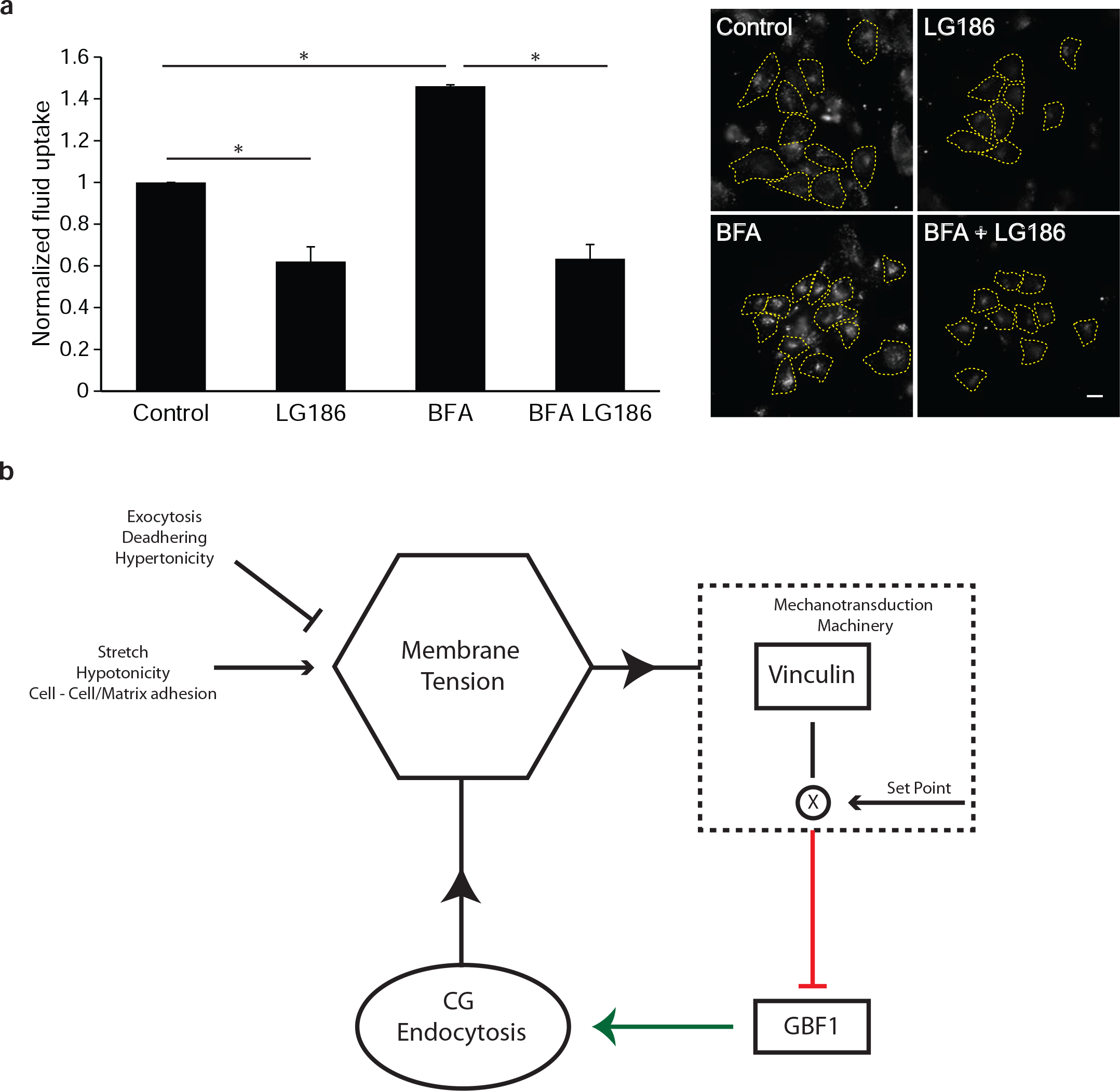
**(a)** Vinculin cells were either untreated (Control) or treated with either LG186 (LG186), BFA (BFA) or BFA followed by LG186 treatment (BFA LG186). Cells were pulsed with F-Dex for 3 minutes and uptake per cell quantified as detailed in Methods, normalized to control, and plotted in the histogram (left). The histogram and Images (right) show that BFA treatment causes an increase in fluid-phase uptake in vinculin null cells and treating with LG186 inhibits fluid-phase uptake even in BFA treated cells. This indicates that the CG pathway is operational and capable of being upregulated in the vinculin null cells. Data represent the mean intensity per cell (± S.D) from two different experiments with duplicates containing at least 100 cells per experiment. *: *P* < 0.001, ns: not significant. Scale bar, 10 μm. **(b)** Model for maintaining membrane tension homeostasis. Different processes such as endocytosis, hypotonic shock, and adhesion increase effective membrane tension (arrow headed line) while exocytosis, hypertonic medium and deadhering would reduce the effective membrane tension (bar headed line). This effective membrane tension is a physical parameter that activates a vinculin dependent mechano-transduction machinery. Depending on the change in tension in relation to a steady state set point, this mechanotransduction machinery inhibits the activity of GBF1 at the cell surface (red bar headed line). Higher effective membrane tension in relation to the steady state would activate vinculin mediated machinery and inhibit GBF1. GBF1 levels at the cell surface is crucial for the activity of the CG pathway and thus directly regulates the level of the endocytosis (green arrow headed line). Thus any modulation of the membrane tension could be balanced by this negative feedback loop to maintain tension homoeostasis.

## METHODS

### Cell Culture and reagents

The CHO (Chinese Hamster Ovary) cells stably expressing FR-GPI and human transferrin receptor TfR (IA2.2 cells) as described before^33^, HeLa, MEF (Mouse embryonic fibroblasts) cells, Caveolin null MEF, conditional null dynamin triple knockout MEF, vinculin null MEF were used for the assays. HF12 media (HiMEDIA, Mumbai, India) and DMEM (Invitrogen) supplemented with NaHCO3 and L-Glutamin/Penicillin/Streptomycin solution (Sigma Aldrich) was used for growing CHO cells and the different MEF lines respectively.

BrefeldinA (BFA) (Sigma Aldrich), ML141 (Tocris Bioscience) and LG186 (see synthesis section below) dissolved in DMSO was used at 20 μg/ml, 10μM and 10μM respectively. ML141 and LG186 treatment was done for 30 minutes in serum free media and maintained during endocytic assays. Tetra methyl rhodamine labelled dextran (TMR-Dex) (10,000MW;Molecular probes, Thermofisher Scientific) was used at 1mg/ml. 4 – hydroxy tamoxifen (Sigma Aldrich) was used at 3 μM to remove Dynamin 1/2/3 from the conditional dynamin triple knockout MEF cells as reported previously ^29^. TrypLE express (GIBCO, Invitrogen) was used to detach cells according to manufacturer’s instruction. SYLGARD 184 silicone elastomer kit (Dow Corning) was used to make PDMS sheets according to manufacturer’s instruction. For the reservoir experiments, cells were transfected with a membrane targeting plasmid pEYFP-mem (Clontech) using the Neon transfection device according to the manufacturer’s protocol as described earlier ^6^.

### Endocytic and Recycling assays on deadhering

Endocytic assays were done as previously described^28,33^ with slight modifications as required. Briefly, CG endocytosis was monitored using fluorescent dextran at 1mg/ml in medium or fluorescent folate analogue (N^α^-pteroyl-N^ɛ^-Bodipy^TMR^-L-lysine (PLB^TMR^)) in folate free medium for indicated time points at 37°C. Endocytosis of TfR was monitored using 10 μg/ml fluorescent transferrin (Tf) at 37°C incubation for indicated time points. Endocytosis was stopped using ice cold HEPES based buffer(M1) (M1:140mM NaCl, 20mM HEPES, 1mM CaCl_2_, 1mM MgCl_2_, 5mM KCl, pH 7.4). To remove surface fluorescence, cells were treated with PI-PLC (50μg/ml, 1h; GPI-APs) or with ascorbate buffer (160mM sodium ascorbate, 40mM ascorbic acid, 1mM MgCl_2_, 1mM CaCl_2_, pH 4.5; Tf) at 4°C and subsequently fixed with 4% paraformaldehyde for 10 minutes.

To study endocytosis on deadhering, cells were detached using TrypLE containing fluorescent dextran at 1mg/ml concentration for 3 minutes and the detached cells were pipetted into an ice cold vial containing M1 buffer to stop the endocytosis. Cells were then replated back on coverslip bottom dish maintained at 4°C, fixed, washed and imaged. To look at endocytosis in suspension, the cells soon after detaching were pipetted into vial containing fluorescent dextran kept at 37°C. The volume was adjusted to have a final concentration of 1mg/ml of the TMR-Dex and after 3minutes the endocytosis was stopped by shifting vial to ice. The cells are spun down at 4°C and then re-plated on coverslip bottom dish coated with ConA, maintained at 4°C, fixed, washed and imaged.

To understand recycling of cargo on deadhering, cells were pulsed with F-Dex for 3 minutes, quickly washed with M1 buffer at room temperature and then detached with TrypLE at 37°C for 5minutes, pipetted into vial containing ice cold M1 buffer and kept on ice. Cells were then re-plated back on coverslip bottom dish coated with ConA maintained at 4°C, fixed, washed and imaged.

For different small molecule inhibitors used, the cells were treated with them for 30 minutes in serum free media in their respective final concentrations and then medium was removed and pulsed with F-Dex at 1mg/ml in serum free media containing the inhibitors since the inhibitor activity is reversible. Endocytosis is stopped by washing with ice cold M1 buffer, fixed and imaged.

### CTxB-HRP uptake, DAB reaction and Electron Microscopy

WT and Cav^−/−^ MEFs were de-adhered at room temperature followed by internalization of 4 μg.ml^−1^ CTxB-HRP (Invitrogen) at 37°C for 5 minutes, washed two times with ice cold PBS followed by incubation on ice for 10 minutes with 1mg.ml^−1^ DAB(Sigma-Aldrich) with 50μM Ascorbic acid. This is followed by a 10 minute treatment with DAB, Ascorbic acid and 0.012% H_2_O_2_ and then washed twice with ice cold PBS. Cells were fixed using 2.5% Glutaraldehyde (ProSciTech) at room temperature (RT) for 1 hour followed by PBS wash for two times and then washed with 0.1M Na cacodylate and left in the same for overnight at 4°C. Cells were contrasted with 1% osmium tetroxide and 4% uranyl acetate. Cells were dehydrated in successive washes of 70%, 90% and 100% ethanol before embedding using 100% LX-112 resin at 60°C overnight. Sections were viewed under a transmission electron microscope (JEOL 1011; JEOL Ltd. Tokyo, Japan), and electron micrographs were captured with a digital camera (Morada; Olympus) using AnalySIS software (Olympus).

### Preparation of PDMS membrane ring

Sylgard 184 silicone elastomer kit comes in two parts which are added in 10 to 1 mix ratio between the polydimethylsiloxane base and the curing agent. This is thoroughly mixed and degassing is done either using a vacuum desiccator or by centrifugation. To prepare PDMS sheets, 7ml of this mixture was added to the middle of a circular 6inch plate which is spun at 500R.P.M for a minute on a spin coater. This was cured at 65°C overnight and then carefully peeled off either after treatment with oxygen plasma cleaner for 40 seconds or without treatment. This PDMS sheet is spread evenly and tightly placed between rings of the stretcher (Fig 1a). The cells were plated in the middle of the PDMS sheet surrounded with water soaked tissue paper to retain humidity and prevent drying up of medium. These rings were placed in the stretcher and stretched by varying the level of vacuum as needed according to the calibration for the experiments.

### Stretch and osmolarity experiments

For the stretch relax experiments, cells plated on PDMS membrane were loaded on the cell stretcher system (Fig 1a) within a temperature controlled chamber at 37°C. Vacuum was applied beneath the ring containing the PDMS sheet, deforming the membrane and stretching the cells plated on the PDMS. The setup was calibrated to stretch cells equi-biaxially to cause 6% strain. Medium containing F-Dex was kept at 37°C in a water bath, and used for the endocytic pulse for the indicated time during or after stretch (see endocytic protocol above). Control cells were treated in the same way except for application of stretch.

For the osmolarity experiments, cells were treated with 50% hypotonic medium made with deionized water at 37°C for the indicated time points and then pulsed with F-Dex either in hypotonic or isotonic medium as needed. The shock is applied for 60 seconds and pulse done for 60 seconds as mentioned. Endocytosis was stopped with ice cold M1 buffer, washed, fixed and imaged.

### Optical Tweezer measurements

Tether forces was measured using an custom built optical tweezer using IR laser (CW,1064nm, TEM_00,1W) along with 100x, 1.3NA oil objective and motorized stage on a Olympus IX71 inverted microscope. Polystyrene beads added to the imaging chamber were allowed to settle and then held in the optical trap while simultaneously imaging through bright field on a coolsnap HQ CCD camera. Membrane tethers are formed by attaching the beads to the cells for few seconds and by moving the bead away using the piezo stage. The tether is held at a constant length and the fluctuation in the trapped bead is detected by using a quadrant photodiode which in turn is acquired and saved using a Labview program through a Data Acquisition Card (USB-6009 NI). The trap stiffness is calibrated using the power spectrum method. The displacement of the bead from the center along with the trap stiffness is used to calculate the tether forces live using a custom written Labview code.

### Magnetic Tweezer measurements

Magnetic tweezer measurements were carried out as previously described^59^. Briefly, a custom-built electromagnet with a sharpened tip was used to apply a 0.5 nN pulsatory force (1 Hz) to paramagnetic beads attached to cells. Paramagnetic beads had been previously coated with either the fibronectin fragment FN7-10 or concanavalin A (Sigma Aldrich), which bind respectively specifically to integrins and the actin cytoskeleton or nonspecifically to the membrane through glycoproteins^60^. Bead movement in response to applied force was then tracked, and bead stiffness was calculated during the first 10 seconds of the measurement as the transfer function between the applied force and the resulting displacement, evaluated at 1 Hz. This stiffness thus provides a measure of the resistance to force of the cell-bead adhesion in units of nN/μm.

### Synthesis of LG186

LG186 was synthesized freshly prior to experiments as reported previously ^36^ with slight modification as described below and used at 10μM dissolved in DMSO․.

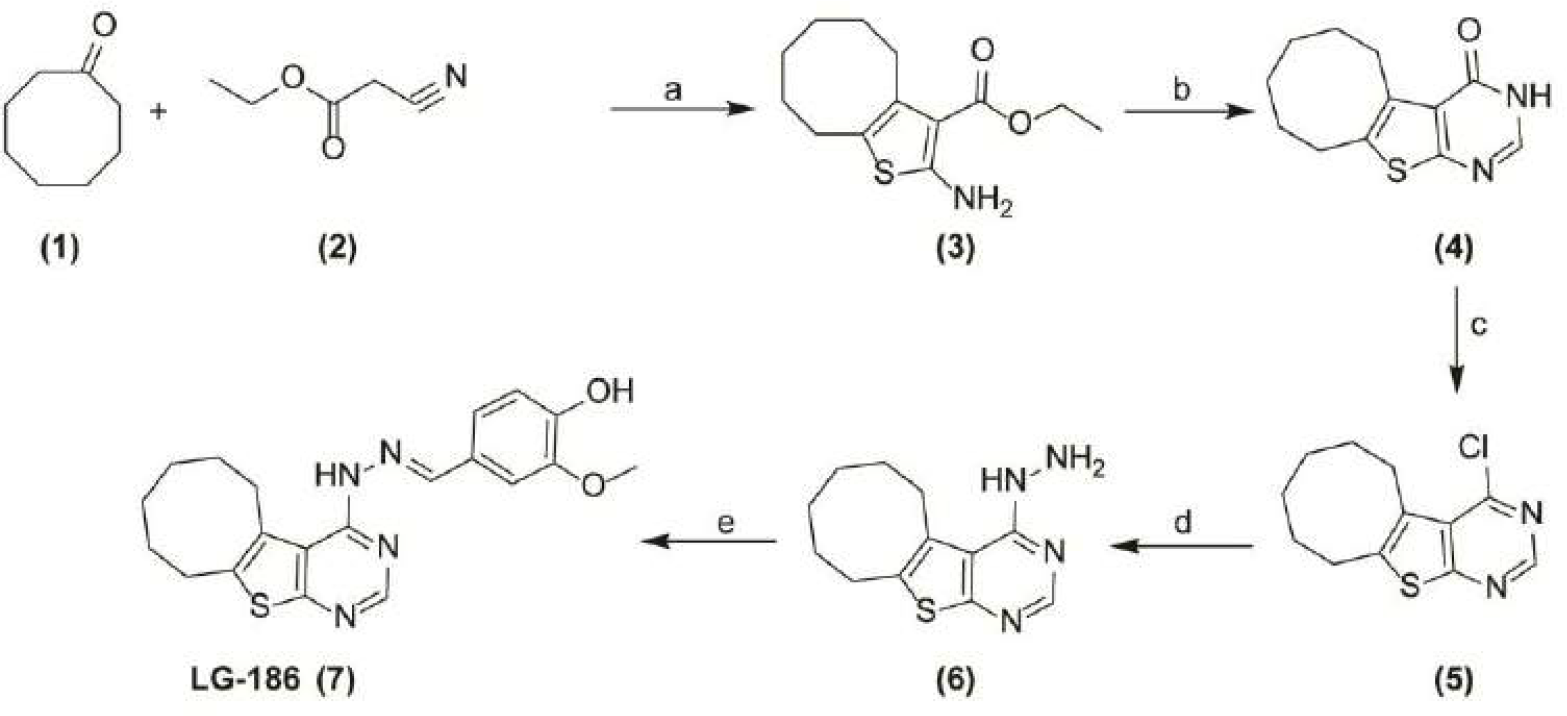

#### Reagent and conditions

Reagents and compounds obtained are mentioned as numbers and the conditions for the reaction are labelled alphabetically along with arrows in figure. a) S_8_, morpholine, ethanol reflux 6 h; b) HCONH_2_, 150°C, 5 h; c) POCl_3_, DMF, rt. d) Hydrazine hydrate, methanol, rt. 2 h; e) Vanillin, rt. 2 h.

#### Compound 3

To a solution of cyclooctanone **(1)** (10 mmol) in ethanol (10 mL) were added sulfur (10 mmol), ethyl cyanoacetate **(2)** (10 mmol) and morpholine (4 mmol). The reaction mixture was stirred at 60°C for 5 h. Upon completion of reaction (checked by TLC), evaporate the solvent and extracted with ethyl acetate and purified by column chromatography using dichloromethane. ^1^H NMR (400 MHz, CDCL_3_) δ: 1.28 (m, 5H), 1.39 (m, 2H), 1.50 (m, 2H), 1.56 (m, 2H), 2.54 (m, 2H), 2.75 (m, 2H), 4.21 (q, 2H), 5.86 (brs, 2H). LC-MS: 254 (M+H)^+^.

#### Compound 4

Compound **3** was heated at 150°C in 5 mL formamide for 5 h. Upon cooling overnight, the product crystallized as slightly brownish crystals. The resulting crystals were collected and washed with a mixture of cold ethanol/water (1/1) to give the corresponding product in quantitative yield. ^1^H NMR (400MHz, DMSO) δ: 1.27(m, 2H), 1.42 (m, 2H), 1.62(m, 4H), 2.87 (m, 2H), 3.06 (m, 2H), 8.5 (s, 1H), 11.4 (brs, 1H). LC-MS: 234.9 (M+H) ^+^.

#### Compound 5

Compound **4** was dissolved in hot DMF and then ice-cooled prior to the addition of POCl_3_ (2 equivalents). Upon stirring overnight, the product precipitated out as white solid **5**, which was collected and washed with cold water and used for next step without purification.

#### Compound 7

Compound **5** was dissolved in methanol and then added 10 equivalent of hydrazine monohydrate. The mixture was stirred for 2 h and water was added. The resulting precipitate was filtered off and washed with cold water to obtain compound **6**, which was then treated with 1.2 equivalent of vanillin. The mixture was stirred for 2 h, diluted with water and extracted with dichloromethane. The organic layer was dried with MgSO4, filtered off and concentrated in vacuo and purified by column chromatography using ethyl acetate: hexane (4:6). ^1^H NMR (400MHz, DMSO) δ: 1.27 (m, 2H), 1.46 (m, 2H), 1.62 (m, 2H), 1.68 (m, 2H), 2.85 (m, 2H), 3.19 (m, 2H), 3.88 (s, 3H), 6.84 (d, 1H), 7.60 (d, 1H), 7.79 (s, 1H), 8.30 (s, 1H), 9.45 (brs, 1H) 11.70 (brs, 1H).^13^C NMR: ^13^C NMR (100 MHz, DMSO) δ 149.29, 148.97, 148.47, 146.76, 146.41, 141.11, 135.01, 132.59, 122.39, 118.77, 117.05, 116.46, 113.05, 58.20, 56.24, 26.95, 25.84, 25.72, 25.50. LC-MS: 383 (M+H)^+^.

### Imaging, Analysis and Statistics

The quantification of endocytic uptake for a population is done by imaging on 20x, 0.75 NA on a Nikon TE300 wide field inverted microscope. For the stretch experiments, an upright microscope (Nikon eclipse Ni −U) was used with a water immersion objective (60x, 1.0 NA). Different fields were obtained and cells within the field were outlined as regions using either metamorph or micromanager software. The images were analyzed using MetaMorph^®^ or Micro-Manager software and is processed for presentation using Adobe Illustrator. All images displayed are equally scaled for intensity unless otherwise mentioned. The scale bar is 10μm unless otherwise mentioned. The integrated intensities, spread area and thus average uptake per cell were determined using the region measurement option. Each value plotted here is mean value from two different experiments with duplicates in each and standard deviation between the two experiments. Statistical significance was tested using Mann-Whitney test and p-values are reported.

